# Morphological plant modeling: Unleashing geometric and topological potential within the plant sciences

**DOI:** 10.1101/078832

**Authors:** Alexander Bucksch, Acheampong Atta-Boateng, Akomian Fortuné Azihou, Mathilde Balduzzi, Dorjsuren Battogtokh, Aly Baumgartner, Brad M. Binder, Siobhan A. Braybrook, Cynthia Chang, Viktoiriya Coneva, Thomas J. DeWitt, Alexander G. Fletcher, Malia A. Gehan, Diego Hernan Diaz Martinez, Lilan Hong, Anjali S. Iyer-Pascuzzi, Laura L. Klein, Samuel Leiboff, Mao Li, Jonathan P. Lynch, Alexis Maizel, Julin N. Maloof, R.J. Cody Markelz, Ciera C. Martinez, Laura A. Miller, Washington Mio, Wojtek Palubicki, Hendrik Poorter, Christophe Pradal, Charles A. Price, Eetu Puttonen, John Reese, Rubén Rellán-Álvarez, Edgar P. Spalding, Erin E. Sparks, Christopher N. Topp, Joseph Williams, Daniel H. Chitwood

## Abstract

Plant morphology is inherently mathematical in that morphology describes plant form and architecture with geometrical and topological descriptors. The geometries and topologies of leaves, flowers, roots, shoots and their spatial arrangements have fascinated plant biologists and mathematicians alike. Beyond providing aesthetic inspiration, quantifying plant morphology has become pressing in an era of climate change and a growing human population. Modifying plant morphology, through molecular biology and breeding, aided by a mathematical perspective, is critical to improving agriculture, and the monitoring of ecosystems with fewer natural resources. In this white paper, we begin with an overview of the mathematical models applied to quantify patterning in plants. We then explore fundamental challenges that remain unanswered concerning plant morphology, from the barriers preventing the prediction of phenotype from genotype to modeling the movement of leafs in air streams. We end with a discussion concerning the incorporation of plant morphology into educational programs. This strategy focuses on synthesizing biological and mathematical approaches and ways to facilitate research advances through outreach, cross-disciplinary training, and open science. This white paper arose from bringing mathematicians and biologists together at the National Institute for Mathematical and Biological Synthesis (NIMBioS) workshop titled “Morphological Plant Modeling: Unleashing Geometric and Topological Potential within the Plant Sciences” held at the University of Tennessee, Knoxville in September, 2015. Never has the need to quantify plant morphology been more imperative. Unleashing the potential of geometric and topological approaches in the plant sciences promises to transform our understanding of both plants and mathematics.

## I. Introduction

### A. Morphology from the perspective of plant biology

The study of plant morphology interfaces with all levels of biological organization **(Figure 1)**. Plant morphology can be descriptive and categorical, as in systematics, which focuses on biological homologies to discern groups of organisms (Mayr, 1981; Wiens, 2000). In plant ecology, the morphology of communities defines vegetation types and biomes, including their relationship to the environment. In turn, plant morphologies are mutually informed by other fields of study, such as plant physiology, the study of the functions of plants, plant genetics, the description of inheritance, and molecular biology, the underlying gene regulation (Kaplan, 2001).

**Figure 1:**
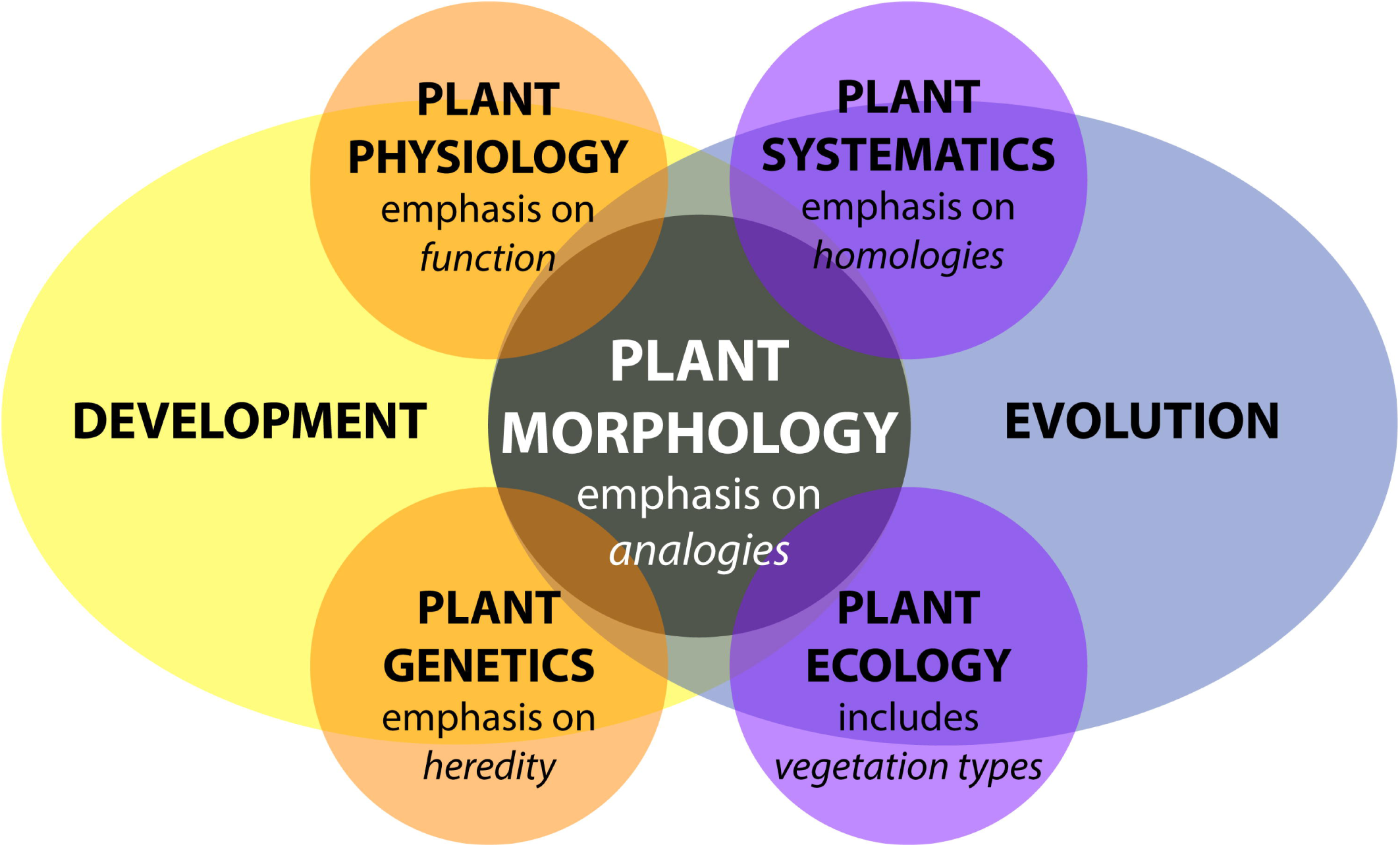
Plant morphology from the perspective of biology. Adapted from Kaplan (2001). Plant morphology interfaces with all disciplines of plant biology—plant physiology, plant genetics, plant systematics, and plant ecology—influenced by both developmental and evolutionary forces.

Plant morphology is more than an attribute affecting plant organization, it is also dynamic. Developmentally, morphology reveals itself over the lifetime of a plant through varying rates of cell division, cell expansion, and anisotropic growth (Esau, 1960; Steeves and Sussex, 1989; Niklas, 1994). Response to changes in environmental conditions further modulate the abovementioned parameters. Development is genetically programmed and driven by biochemical processes that are responsible for physical forces that change the observed patterning and growth of organs (Green, 1999; Peaucelle et al., 2011; Braybrook and Jönsson, 2016). In addition, external physical forces affect plant development, such as heterogeneous soil densities altering root growth or flows of air, water, or gravity modulating the bending of branches and leaves (Moulia & Fournier, 2009). Inherited modifications of structure or development, either incrementally or abruptly, over generations results in the evolution of plant morphology (Niklas, 1997). A record of these changes over geologic time is preserved through fossils and correlates with the paleoclimate, which contributes to our understanding of morphology in extant plants today (Bailey and Sinnott, 1915). Development and evolution are the biological mechanisms through which plant morphology arises, regardless of whether in a systematic, ecological, physiological, or genetic context **(Figure 1)**.

### B. Plant morphology from the perspective of mathematics

In 1790 Johann Wolfgang von Goethe pioneered a perspective that transformed the way mathematicians think about plant morphology: the idea that the essence of plant morphology is an underlying repetitive process of deformation (Goethe, 1790; Friedman and Diggle, 2011). The modern challenge that Goethe’s paradigm presents is to quantitatively describe deformations resulting from differences in the underlying genetic, developmental, and environmental cues. From a mathematical perspective, the challenge is how to define shape descriptors to compare, and generation processes to simulate, plant morphology with topological and geometrical techniques.

#### 1. Mathematics to describe plant shape and morphology

Several areas of mathematics can be used to extract quantitative measures of plant shape and morphology. One intuitive representation of the plant form relies on the use of skeletal descriptors that reduce the branching morphology of plants to a set of intersecting lines or curve segments, constituting a mathematical graph. These skeleton-based mathematical graphs can be derived from manual measurement (Godin et al., 1999; Watanabe et al., 2005) or imaging data (Bucksch et al., 2010; Bucksch 2011; Bucksch, 2014a; Aiteanu and Klein, 2014). Such skeletal descriptions can be used to derive quantitative measurements of lengths, diameters, and angles in tree crowns (Bucksch and Fleck, 2011; Raumonen et al., 2013; Seidel et al., 2015) and roots, at a single time point (Fitter, 1987; Danjon et al., 1999; Lobet et al., 2011; Galkovskyi et al., 2012) or over time to capture growth dynamics (Symonova et al., 2015). Having a skeletal description in place allows the definition of orders, in a biological and mathematical sense, to enable morphological analysis from a topological perspective **(Figure 2A)**. Topological analyses can be used to compare shape characteristics independently of events that deform and transform plant shape geometrically, providing a framework by which plant morphology can be modeled. The relationships between orders, such as degree of self-similarity (Prusinkiewicz, 2004) or self-nestedness (Godin and Ferraro, 2010) are used to quantitatively summarize patterns of plant morphology. Persistent homology **(Figure 2B)**, an extension of Morse theory (Milnor, 1963), deforms a given plant shape gradually to define self-similarity (MacPherson and Schweinhardt, 2012) and morphological properties (Edelsbrunner and Harer, 2010) on the basis of topological event statistics. In the example in **Figure 2B**, topological events are represented by the geodesic distance at which branches are “born” and “die” along the length of the structure.

**Figure 2:**
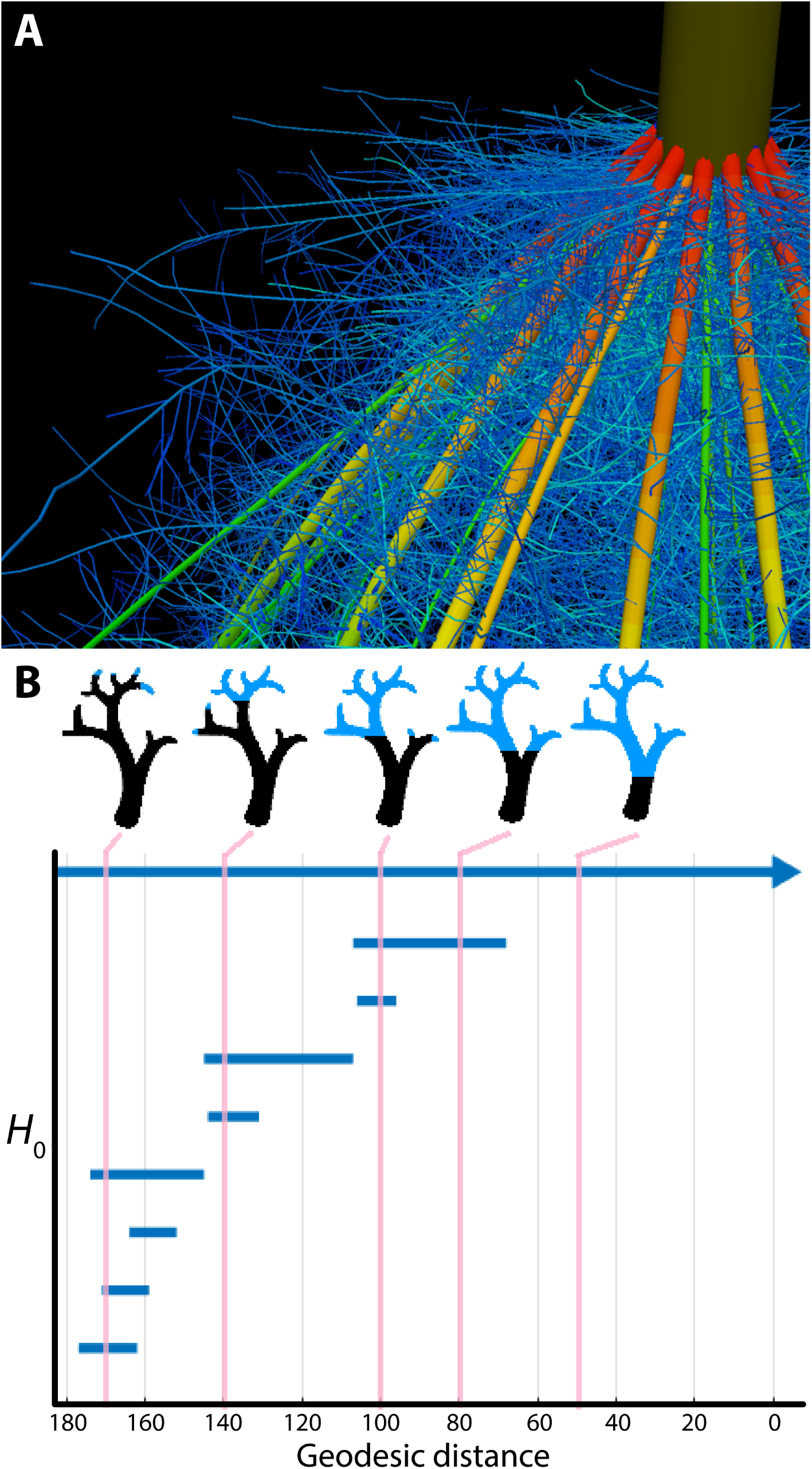
Plant morphology from the perspective of mathematics. A) The topological complexity of plants requires a mathematical framework to describe and simulate plant morphology. Shown is the top of a maize crown root 42 days after planting. Color represents root diameter, revealing topology and different orders of root architecture. Image provided by Jonathan Lynch and Johannes Postma (Pennsylvania State University). B) Persistent homology deforms a given plant morphology using functions to define self-similarity in a structure. In this example, a geodesic distance function is traversed to the ground level of a tree (that is, the shortest curved distance of each voxel to the base of the tree), as visualized in blue in successive images. The branching structure, as defined across scales of the geodesic distance function is recorded as an *H*_0_ (zero-order homology) barcode, which in persistent homology refers to connected components. As the branching structure is traversed by the function, connected components are “born” and “die” as terminal branches emerge and fuse together. Each of these components is indicated as a bar in the *H*_0_ barcode, and the correspondence of the barcode to different points in the function is indicated by vertical lines, in pink. Images provided by Mao Li (Danforth Plant Science Center).

Traditionally, descriptors that compare outlines of plant organs independently of scale, rotation, and translation have been used to quantify morphologies. However, in the 1980s, David Kendall defined an elegant alternative statistical framework to these descriptors (Kendall, 1984). His idea was to compare the outline of shapes in a transformation-invariant fashion, which fulfills the parameters of the mathematical concept of shape. This concept infused rapidly as morphometrics into biology (Bookstein, 1997) and is increasingly carried out using machine vision techniques (Wilf et al., 2016). Kendall’s idea inspired the development of methods such as elliptical Fourier descriptors (Kuhl and Giardina, 1982) and new trends employing the Laplace Beltrami operator (Reuter et al., 2009), both relying on the spectral decompositions of shapes (Chitwood et al., 2012a; Chitwood et al., 2012b; Laga et al. 2014; Rellán-Álvarez et al., 2015). Beyond the organ level, such morphometric descriptors were used to analyze cellular expansion rates of rapidly deforming primordia into mature organ morphologies (Rolland-Lagan et al., 2003; Remmler and Rolland-Lagan, 2012; Das Gupta and Nath, 2015).

Parallel to strictly mathematical descriptions of plant morphology, Ronald Fisher developed a statistical framework to partition variance into different sources of variability (Fisher, 1925). Specifically, with respect to plant morphology, the *Iris* flower dataset (Fisher, 1936) was used to develop novel methods to differentiate three *Iris* species based on the length and width of sepals and petals.

From a geometric perspective, developmental processes construct surfaces in a three-dimensional space. Yet, this space in which development is embedded imposes constraints on plant forms observed. Awareness of these constraints has led to new interpretations of plant morphology (Prusinkiewicz and DeReuille, 2010; Bucksch et al., 2014b) that might provide avenues to explain symmetry or asymmetry in leaf shape (Martinez et al., 2016) or the occurrence of plasticity in leaf shape as the morphological response of plants to environmental changes over developmental and evolutionary timescales (Royer et al., 2009; Chitwood et al., 2016).

#### 2. Mathematics to simulate plant morphology

Computer simulations use principles from graph theory, such as graph rewriting, to model plant morphology over developmental time by successively augmenting a graph with vertices and edges as plant development unfolds using observed rules (Hallé, 1971). These rules unravel the differences between observed plant morphologies across plant species (Kurth, 1994; Prusinkiewicz et al., 2001; Barthélémy and Caraglio, 2007) and are capable of modeling fractal descriptions that reflect the repetitive and modular appearance of branching structures (Horn, 1971, Hallé, 1986). Recent developments in graph theory abstract the genetic mechanisms driving the developmental program of tree crown morphology into a computational framework (Runions et al., 2007; Palubicki et al., 2009; Palubicki, 2013). Equivalently, functional-structural models of roots can be utilized to simulate the efficiency of nutrient and water uptake following developmental programs (Nielsen et al., 1994; Dunabin et al., 2013).

Alan Turing, a pioneering figure in twentieth-century science, had a longstanding interest in phyllotactic patterns. Turing’s approach to the problem was twofold: first, a detailed geometrical analysis of the patterns (Turing, 1992), and second, an application of his theory of morphogenesis through local activation and long-range inhibition (Turing, 1952), which defined the first reaction-diffusion system for morphological modeling. Combining physical experiments with computer simulations, Douady and Coudert (1996) subsequently modeled a diffusible chemical produced by a developing primordium that would inhibit the initiation of nearby primordia, successfully recapitulating known phyllotactic patterns in the shoot apical meristem (Bernasconi, 1994; Meinhardt, 2004; Hohm et al., 2010; Fujita et al., 2011), the number of floral organs (Kitazawa and Fujimoto, 2015), the regular spacing of root hairs (Meinhardt and Gierer, 1974), and the establishment of specific vascular patterns (Meinhardt, 1976).

## II. Emerging questions and barriers in the mathematical analysis of plant morphology

A true synthesis of plant morphology, which comprehensively models observed biological phenomena and incorporates a mathematical perspective, remains elusive. In this section we highlight current focuses in the study of plant morphology, including the technical limits of acquiring morphological data, phenotype prediction, responses of plants to the environment, models across biological scales, and the integration of complex phenomena, such as fluid dynamics, into plant morphological models.

### A. Technological limits acquiring plant morphological data

There are several technological limits to acquiring plant morphological data that must be overcome to move this field forward. One such limitation is the acquisition of quantitative plant images. Traditionally, many acquisition systems do not provide morphological data with measurable units. Approaches that rely on the reflection of waves from the plant surface can provide quantitative measurements for morphological analyses. Time of flight scanners, such as terrestrial laser scanning, overcome unit-less measurement systems by recording the round-trip time of hundreds of thousands of laser beams sent at different angles from the scanner to the first plant surface within the line of sight (Vosselman and Maas, 2010) **(Figure 3)**. Leveraging the speed of light allows calculation of the distance between a point on the plant surface and the laser scanner. Laser scanning and the complementary approach of stereovision both produce surface samples or point clouds as output. However, both approaches face algorithmic challenges encountered when plant parts occlude each other, since both rely on the reflection of waves from the plant surface (Bucksch, 2014a). Radar provides another non-invasive technique to study individual tree and forest structures over wide areas. Radar pulses can either penetrate or reflect from foliage, depending on the selected wavelength (Kaasalainen et al., 2015). Most radar applications occur in forestry and are being operated from satellites or airplanes. Although more compact and agile systems are being developed for precision forestry above- and below-ground (Feng et al., 2016), their resolution is too low to acquire the detail in morphology needed to apply hierarchy or similarity oriented mathematical analysis strategies.

**Figure 3:**
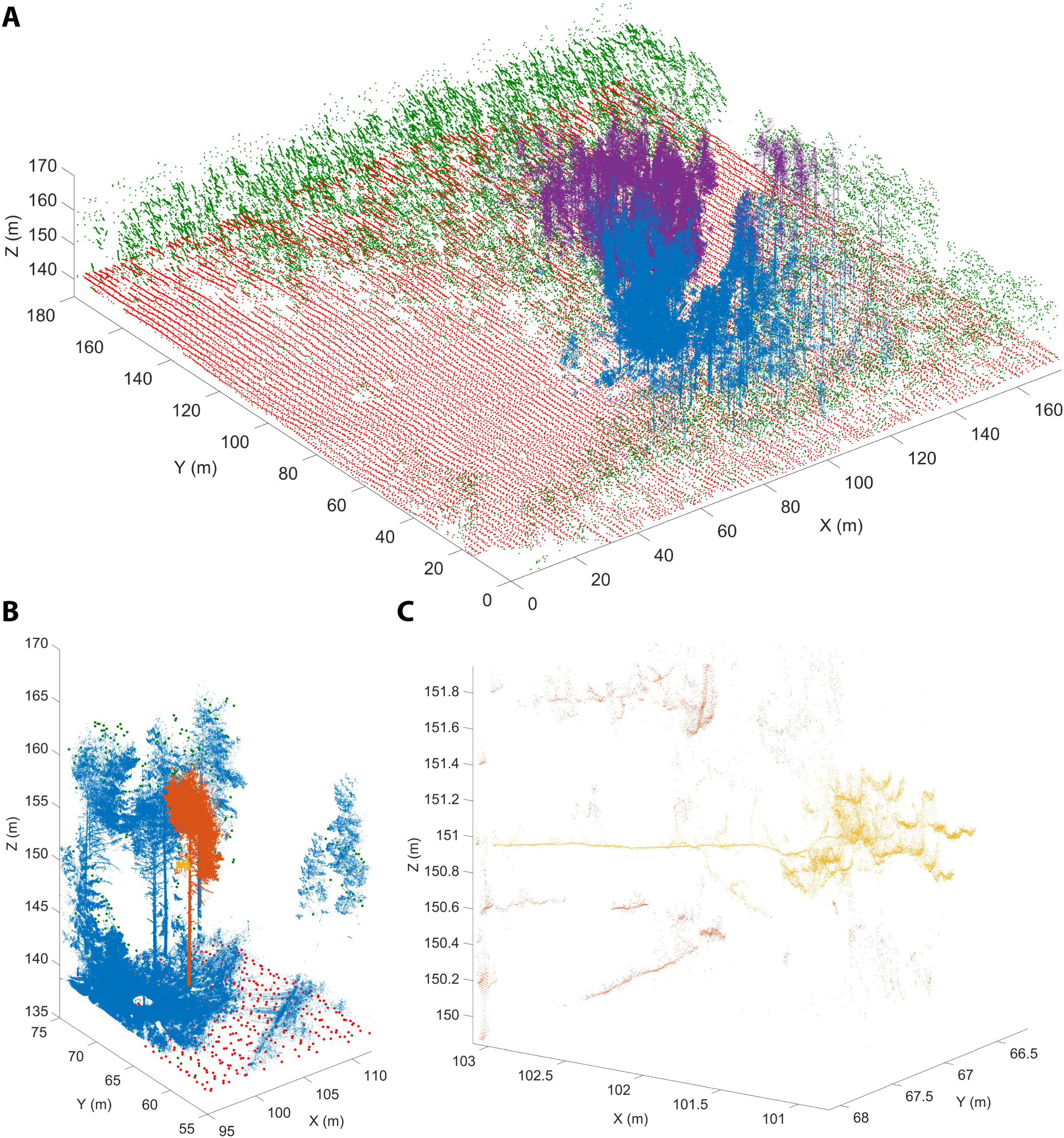
Terrestrial laser scanning creates a point cloud reconstruction of a Finnish forest. A) Structure of a boreal forest site in Finland as seen with airborne (ALS) and terrestrial (TLS) laser scanning point clouds. The red (ground) and green (above-ground) points are obtained from National Land Survey of Finland national ALS point clouds that cover hundreds of thousands of square kilometers with about 1 point per square meter resolution. The blue and magenta point clouds are results of two individual TLS measurements and have over 20 million points each within an area of about 500 square meters. TLS point density varies with range but can be thousands of points per square meter up to tens of meters away from the scanner position. B) An excerpt from a single TLS point cloud (blue). The TLS point cloud is so dense that individual tree point clouds (orange) and parts from them (yellow) can be selected for detailed analysis. C) A detail from a single TLS point cloud. Individual branches (yellow) 20 meters above ground can be inspected from the point cloud with centimeter level resolution to estimate their length and thickness. Images provided by Eetu Puttonen (Finnish Geospatial Research Institute in the National Land Survey of Finland). ALS data was obtained from the National Land Survey of Finland Topographic Database, 08/2012 (National Land Survey of Finland open data licence, version 1.0).

Image techniques that utilize penetration of the plant tissue to resolve occlusions are possible with X-ray (Kumi et al., 2015) and magnetic resonance imaging (MRI; van Dusschoten et al., 2016). While both technologies resolve occlusions and can even penetrate soil, their limitation is the requirement of a closed imaging volume. Thus, although useful for a wide array of purposes, MRI and X-ray are potentially destructive if applied to mature plant organs such as roots in the field or tree crowns that are larger than the imaging volume (Fiorani et al., 2012). Interior plant anatomy can be imaged using confocal microscopy and laser ablation **(Figure 4)** or nano- or micro-CT tomography techniques, that are limited to small pot volumes, to investigate the first days of plant growth.

**Figure 4:**
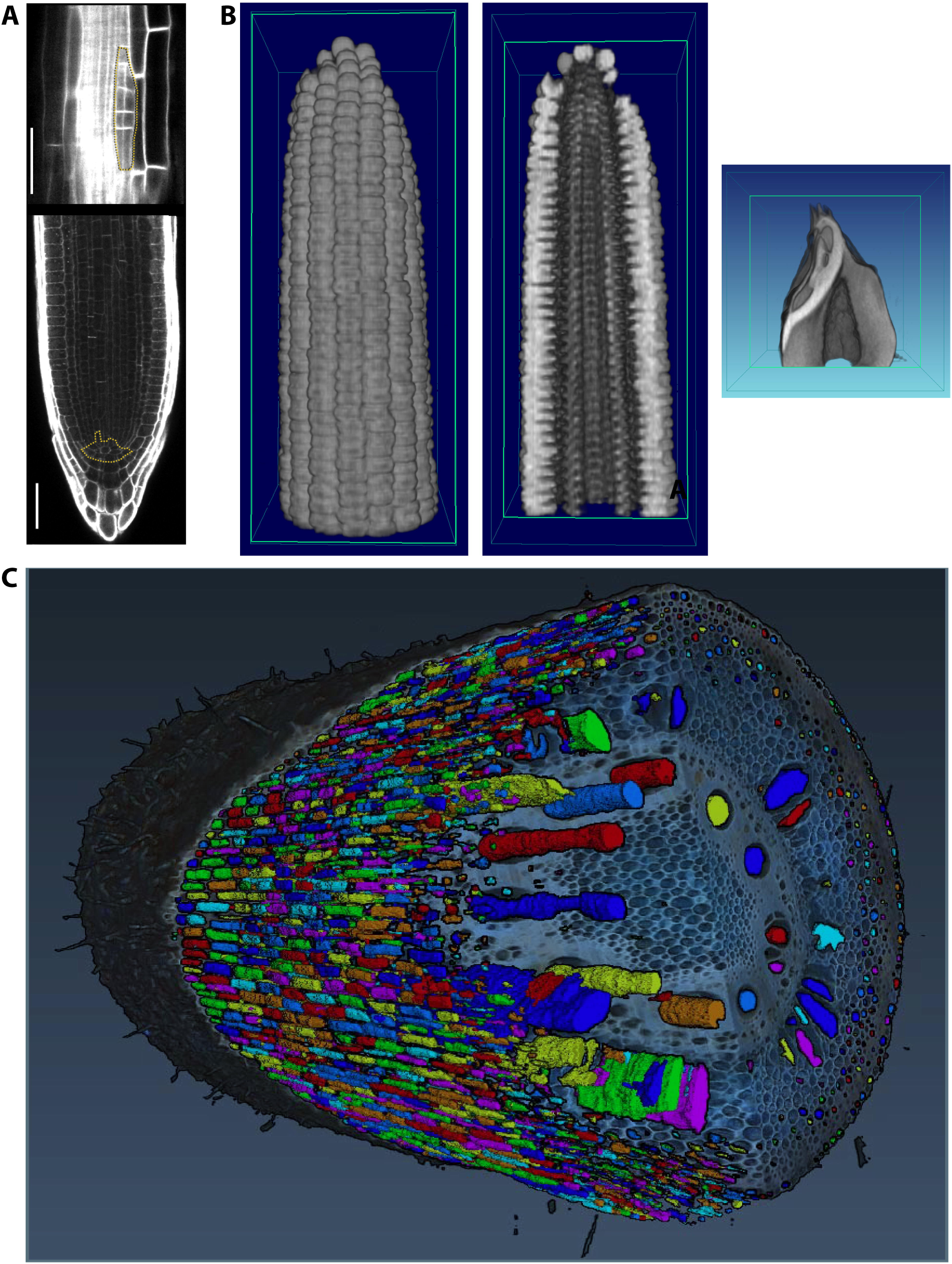
Imaging techniques to capture plant morphology. A) Confocal sections of an Arabidopsis root. The upper panel shows a new lateral root primordium at an early stage of development (highlighted in yellow). At regular intervals new roots branch from the primary root. The lower panel shows the primary root meristem and the stem cell niche (highlighted in yellow) from which all cells derive. Scale bars: 100µm. Images provided by Alexis Maizel (Heidelberg University). B) Computational tomographic (CT) x-ray sections through a reconstructed maize ear (left and middle) and kernel (right). Images provided by Chris Topp (Donald Danforth Plant Science Center). C) Laser ablation tomography (LAT) image of a nodal root from a mature, field-grown maize plant, with color segmentation showing definition of cortical cells, aeraenchyma lacunae, and metaxylem vessels. Image provided by Jennifer Yang (Penn State).

### B. The genetic basis of plant morphology

One of the outstanding challenges in plant biology is to link the inheritance and activity of genes with observed phenotypes. This is particularly challenging for the study of plant morphology, as both the genetic landscape and morphospaces are complex: modeling each of these phenomena alone is difficult, let alone trying to model morphology as a result of genetic phenomena (Benfey and Mitchell-Olds, 2008; Lynch and Brown, 2012; Chitwood and Topp, 2015). Although classic examples exist in which plant morphology is radically altered by the effects of a few genes (Doebley, 2004; Clark et al., 2006; Kimura et al., 2008), many morphological traits have a polygenic basis (Langlade et al., 2005; Tian et al., 2011; Chitwood et al., 2013; 2014b).

Quantitative trait locus (QTL) analyses can identify the polygenic basis for morphological traits that span scales from the cellular to the whole organ level. At the cellular level, root cortex cell number (Ron et al., 2013), the cellular basis of carpel size (Frary et al., 2000), and epidermal cell area and number (Tisne et al., 2008) have been analyzed. The genetic basis of cellular morphology ultimately affects organ morphology, and quantitative genetic bases for fruit shape (Monforte, et al., 2014; Paran and van der Knaap, 2007), root morphology (Zhu et al., 2005; Clark et al., 2011; Topp et al., 2013; Zurek, et al., 2015), shoot apical meristem shape (Thompson et al., 2015; Leiboff et al., 2015), leaf shape (Langlade et al., 2005; Ku et al., 2010; Tian et al., 2011; Chitwood et al., 2013; 2014a; 2014b; Zhang et al., 2014; Truong et al., 2015), and tree branching (Kenis and Keulemans, 2007; Segura et al., 2009) have been described.

Natural variation in cell, tissue, or organ morphology ultimately impacts plant physiology. For example, root cortical aerenchyma formation reduces the metabolic costs of soil exploration, thereby improving plant growth under conditions of suboptimal availability of water and nutrients (Zhu et al. 2010; Postma and Lynch, 2011; Lynch et al., 2013). Maize genotypes with greater root cortical cell size or reduced root cortical cell file number also have reduced metabolic costs, and therefore root deeper to increase water capture under drought (Chimungu et al., 2015). The radial distribution of auxin in the rice root leads to differential cell expansion and deeper root angles, resulting in greater water capture in soils with retracting water tables (Uga et al., 2013).

High-throughput phenotyping techniques are increasingly used to reveal the genetic basis of natural variation. In doing so, phenotyping techniques complement classic approaches of reverse genetics and often lead to novel insights, even in a well-studied species like *Arabidopsis thaliana*. Phenotyping techniques have revealed a genetic basis for such dynamic traits as root growth (Slovack et al., 2014). Similarly, high-resolution sampling of root gravitropism has led to an unprecedented understanding of the dynamics of the genetic basis of plasticity (Miller et al., 2007; Brooks et al., 2010; Spalding and Miller, 2013).

### C. The environmental basis of plant morphology

Plasticity is defined as the ability of one genotype to produce different phenotypes based on environment (Bradshaw 1965; DeWitt and Scheiner, 2004) and adds to the phenotypic complexity created by genetics and development. Trait variation in response to the environment has been defined classically using reaction norms (originally “Reaktionsnorm”) where the value of a certain trait is plotted against different environments (Woltereck, 1909). If the reaction norm line is flat, the trait is not plastic and can be considered canalized across environments; if the reaction norm varies across the environment the trait is plastic and the slope of the reaction norm line will be a measure of the plasticity. Significant differences in slopes among genotypes indicate a genotype by environment (GxE) interaction (Via and Lande, 1985).

Seminal work by Clausen, Keck, and Hiesey (1941) demonstrated using several clonal species in a series of reciprocal transplants that, although heredity exerts the most measureable effects on plant morphology, environment is also a major source of phenotypic variability. Research continues to explore the range of phenotypic variation expressed by a given genotype in the context of different environments, which has important implications for many fields, including conservation, evolution, and agriculture (Nicotra et al., 2010; DeWitt, 2016). Many studies examine phenotypes across latitudinal or altitudinal gradients, or other environmental clines, to characterize the range of possible variation and its relationship to local adaptation processes (Cordell et al. 1998; Díaz et al., 2016).

Below-ground, plants encounter diverse sources of environmental variability, including water availability, soil chemistry, and physical properties like soil hardness and movement. These factors vary between individual plants (Razak et al., 2013) and within an individual root system, where plants respond at spatio-temporal levels to very different granularity (Drew, 1975; Robbins and Dinneny, 2015). Plasticity at a micro-environmental scale has been linked to developmental and molecular mechanisms (Bao et al., 2014). The scientific challenge here is to integrate these effects at a whole root system level and use different scales of information to understand the optimal acquisition in resource limited conditions (Rellán-Álvarez, et al., 2016) **(Figure 5)**.

**Figure 5:**
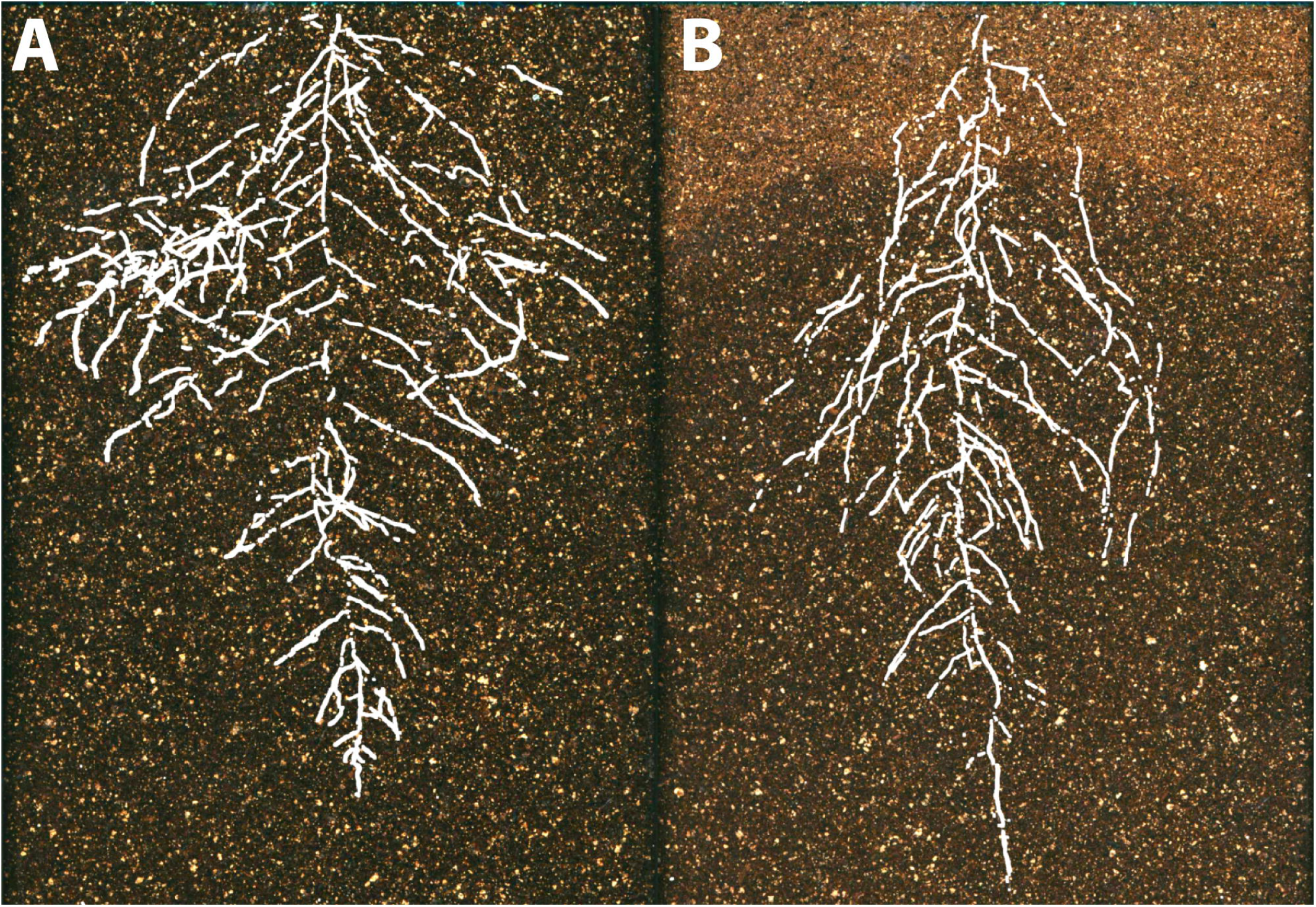
The environmental basis of plant morphology. Root system architecture of Arabidopsis Col-0 plants expressing ProUBQ10: LUC2o growing in A) control and B) water-deficient conditions using the GLO-Roots system (Rellán-Álvarez et al., 2015). Images provided by Ruben Rellán-Álvarez (Laboratorio Nacional de Genómica para la

### D. Integrating models from different levels of organization

Since it is extremely difficult to examine complex interdependent processes occurring at multiple spatio-temporal scales, mathematical modeling can be used as a complementary tool with which to disentangle component processes and investigate how their coupling may lead to emergent patterns at a systems level (Hamant, 2008; Band and King, 2012; Jenzen and Fozard 2015; Band et al. 2012). To be practical, a multiscale model should generate well-constrained predictions despite significant parameter uncertainty (Gutenkunst et al., 2007, Hofhuis et al., 2016). It is desirable that a multiscale model has certain modularity in its design such that individual modules are responsible for modeling specific spatial aspects of the system (Baldazzi et al., 2012). Global sensitivity analysis can be applied to reveal how individual modules function when other modules are perturbed (Sudret, 2008). Most importantly, a multiscale model must be tested against available data (Gordon et al. 2009, Chickarmane et al. 2010, Sahlin et al. 2011, Shapiro et al. 2013, Willis et al. 2016).

To illustrate the challenges of multi-scale modeling, we highlight an example that encompasses molecular and cellular scales. At the molecular scale, models can treat some biomolecules as diffusive, but others, such as membrane-bound receptors, can be spatially restricted (Fujita et al., 2011; Battogtokh and Tyson, 2016). Separately, at the cellular scale, mathematical models describe dynamics of cell networks where the mechanical pressures exerted on the cell walls are important factors for cell growth and division (Jensen and Fozard, 2015) **(Figure 6A)**. In models describing plant development in a two-dimensional cross-section geometry, cells are often modeled as polygons defined by walls between neighboring cells. The spatial position of a vertex, where the cell walls of three neighboring cells coalesce, is a convenient variable for mathematical modeling of the dynamics of cellular networks (Prusinkiewicz and Lindenmayer, 2012). A multiscale model can then be assembled by combining the molecular and cellular models. Mutations and deletions of the genes encoding the biomolecules can be modeled by changing parameters. By inspecting the effects of such modifications on the dynamics of the cellular networks, the relationship between genotypes and phenotypes can be predicted. For example, Fujita et al. (2011) model integrates the dynamics of cell growth and division with the spatiotemporal dynamics of the proteins involved in stem cell regulation and simulates shoot apical meristem development in wild type and mutants plants **(Figure 6B)**.

**Figure 6:**
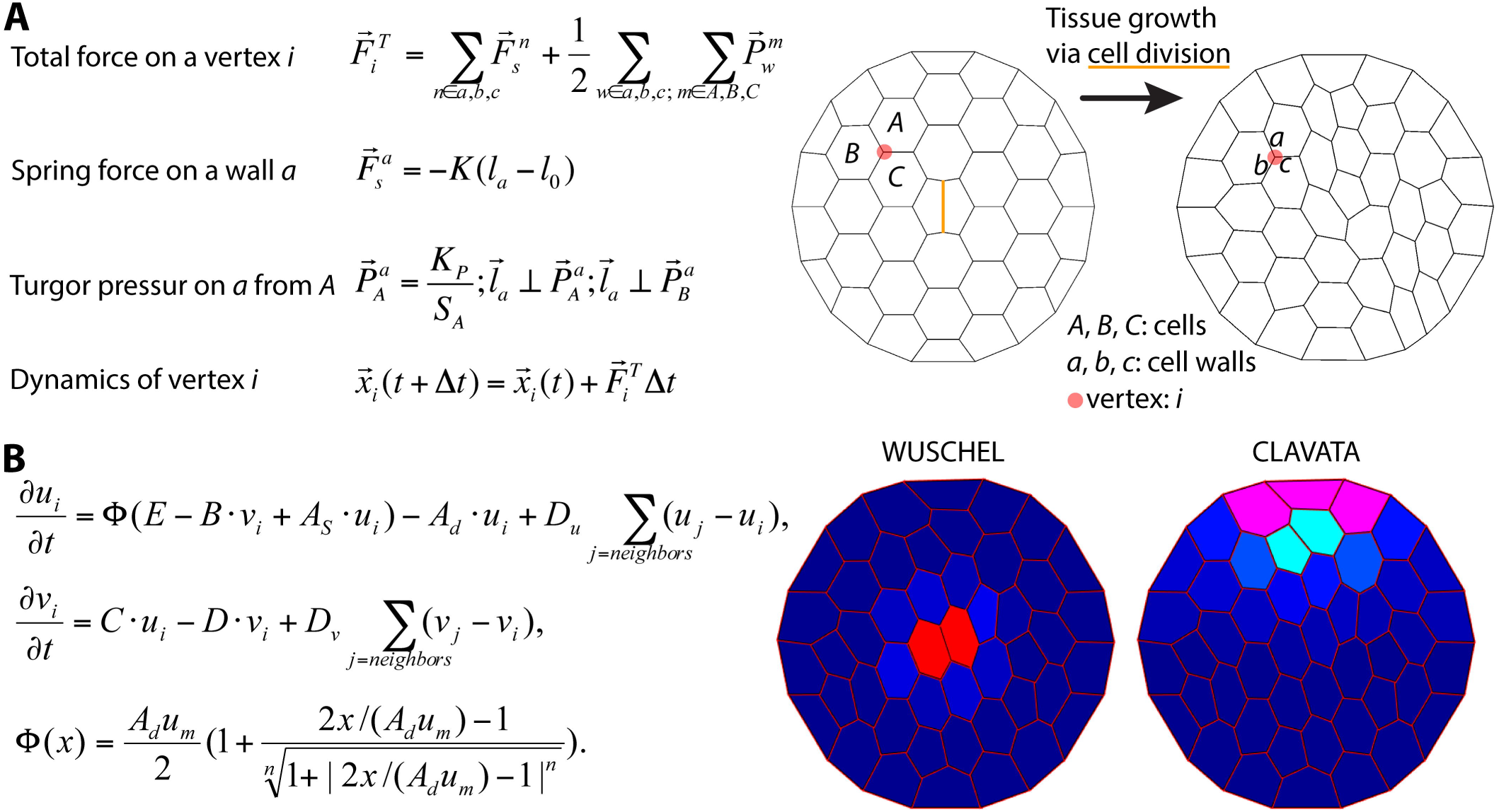
Integration of tissue growth and reaction-diffusion models. A) Vertex model of cellular layers (Prusinjiewicz and Lindenmayer, 2012). K, la, and l0 are the spring constant, current length, and rest length for wall a. KP is a constant and SA is the size of cell A. Δt is time step. Shown is a simulation of cell network growth. B) Reaction diffusion model of the shoot apical meristem for WUSCHEL and CLAVATA interactions (Fujita et al., 2011) u=WUS, v=CLV, i=cell index, Φ is a sigmoid function. E, B, AS, Ad, C, D, um, Du, Dv are positive constants. Shown are the distributions of WUS and CLV levels within a dynamic cell network. Images provided by Dorjsuren Battogtokh (Virginia Tech).

### E. Modeling the impact of morphology on plant function

Quantitative measures of plant morphology are critical to understand function. In one example, leaf shape and material properties that alter the boundary layer of the fluid over the surface of the leaf or enhance passive movement can potentially augment gas and heat exchange. For example, it has been proposed that the broad leaves of some trees flutter for the purpose of convective and evaporative heat transfer (Thom, 1968; Grant, 1983). Fluttering may also allow more light to penetrate the canopy (Roden and Pearcy, 1993).

The morphology and mechanical properties of leaves can alter the boundary layer. For example, trichomes, the hair-like protrusions on the surfaces of leaves, can effectively thicken the boundary layer around a leaf under some conditions (Benz and Martin, 2006) and increase turbulence (Schreuder et al., 2001). Any movement of the leaf relative to the movement of the air or water may, in some cases, act to decrease the boundary layer and increase gas exchange, evaporation, and heat dissipation (Roden and Pearcy, 1993). Each of these parameters may be altered by the plant to improve the overall function of the leaf (Vogel, 2012).

Vogel (1989) was the first to provide quantitative data on drag reduction in plants. He found that single broad leaves reconfigure at high flow velocities into cone shapes that reduce flutter and drag when compared to paper cut-outs of similar shape and flexibility **(Figure 7A-B)**. Subsequent experimental studies on broad leaves, compound leaves, and flowers also support rapid repositioning in response to strong currents as a general mechanism to reduce drag (Vogel, 1989; Niklas, 1992; Ennos, 1997; Etnier and Vogel, 2000; Vogel, 2006) **(Figure 7C)**.

**Figure 7:**
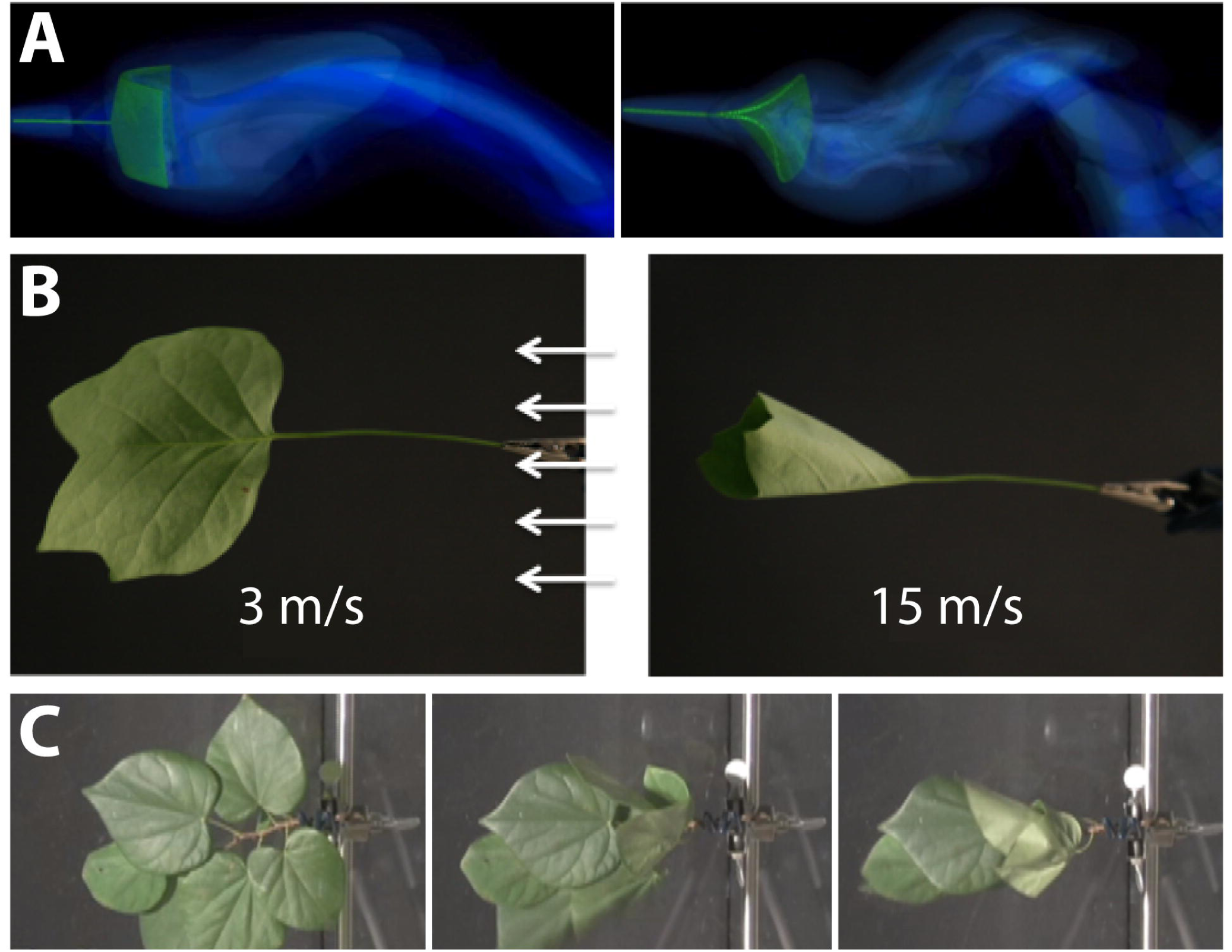
Modeling the interaction between plant morphology and fluid dynamics. A) 3D immersed boundary simulations of flow past a flexible rectangular sheet (left) and disk with a cut from the center to edge (right). Both structures are attached to a flexible petiole, and the flow is from left to right. The contours show the magnitude of vorticity (the rotation in the air). The circular disk reconfigures into a cone shape, similar to many broad leaves. B) Reconfiguration of tulip poplar leaves in 3 m/s (left) and 15 m/s flow (right). The leaves typically flutter at lower wind speeds and reconfigure into stable cones at high wind speeds. C) A cluster of redbud leaves in wind moving from right to left. The wind speed is increased from 3 m/s (left) to 6 m/s (middle) and 12 m/s (right). Note that the entire cluster reconfigures into a cone shape. This is different from the case of tulip poplars and maples where each leaf individually reconfigures into a conic shape. Images provided by Laura Miller (University of North Carolina, Chapel Hill).

## III. Milestones in education and outreach to accelerate the infusion of math into the plant sciences

In a world increasingly geared towards a quantitative mindset and with dwindling natural resources both mathematics and plant biology are timely disciplines. These disciplines need to come together through opportunities to interact, including cross-disciplinary training, workshops, meetings, and funding opportunities. Both fields can immediately benefit from more open approaches to science. In this section, we outline perspectives for enhancing the crossover between mathematics and plant biology.

### A. Education

Mathematics has been likened to “biology’s next microscope”, because of the insights into an otherwise invisible world it has to offer. Conversely, biology has been described as “mathematics’ next physics”, stimulating novel mathematical approaches because of the hitherto unrealized phenomena that biology studies (Cohen, 2004). The scale of the needed interplay between mathematics and plant biology is enormous and may lead to new science disciplines at the interface of both: ranging from the cellular, tissue, organismal, and community levels to the global; touching upon genetic, transcriptional, proteomic, metabolite, and morphological data; studying the dynamic interactions of plants with the environment or the evolution of new forms over geologic time; and spanning quantification, statistics, and mechanistic mathematical models.

Research is becoming increasingly interdisciplinary, and undergraduate, graduate, and post-graduate groups are actively trying to bridge the archaic separation between mathematics and biology skillsets. While many graduate programs have specialization tracks under the umbrella of mathematics or biology-specific programs, more frequently departments are forming specially designed graduate groups for mathematical biology. We emphasize the need for more of these graduate groups and the incorporation of mathematics into biology graduate education. This will necessitate team-teaching across disciplines to train the next generation of mathematical biologists.

### B. Public outreach: Citizen science and the maker movement

Citizen science, which is a method to make the general public aware of scientific problems and employ their help in solving them^1^, is an ideal platform to initiate a synthesis between plant biology and mathematics because of the relatively low cost and accessibility of each field. Arguably, using citizen science to collect plant morphological diversity has already been achieved, but has yet to be fully realized. In total, it is estimated that the herbaria of the world possess greater than 207 million voucher specimens^2^, representing the diverse lineages of land plants collected over their respective biogeographies over a timespan of centuries. Digital documentation of the millions of vouchers held by the world’s botanic gardens is actively underway, allowing for researchers and citizens alike to access and study for themselves the wealth of plant diversity across the globe and centuries (Smith et al., 2003; Corney et al., 2012; Ryan, 2013).

The developmental changes in plants responding to environmental variability and microclimatic changes over the course of a growing season can be analyzed by studying phenology. Citizen science projects such as the USA National Phenology Network^3^ or Earthwatch^4^ and associated programs such as My Tree Tracker^5^ document populations and individual plants over seasons and years, providing a distributed, decentralized network of scientific measurements to study the effects of climate change on plants.

Citizen science is also enabled by low-cost, specialized equipment. Whether programing a camera to automatically take pictures at specific times or automating a watering schedule for a garden, the maker movement—a do-it-yourself cultural phenomenon that intersects with hacker culture—focuses on building custom, programmable hardware, whether via electronics, robotics, 3D-printing, or time-honored skills such as metal- and woodworking. The focus on programming is especially relevant for integrating mathematical approaches with plant science experiments. The low-cost of single-board computers (like Raspberry Pi, Hummingboard, or Cubieboard) makes tinkering more permissive for a greater population of citizen scientists than previously feasible.

### C. Workshops and funding opportunities

Simply bringing mathematicians and plant biologists together to interact, to learn about tools, approaches, and opportunities in each discipline that researchers may not be aware of, is a major opportunity for the full integration of these two disciplines. This white paper itself is a testament to the power of bringing mathematicians and biologists together, resulting from a National Institute for Mathematical and Biological Synthesis (NIMBioS) workshop titled “Morphological Plant Modeling: Unleashing Geometric and Topologic Potential within the Plant Sciences”, held at the University of Tennessee, Knoxville September 2-4, 2015^6^ **(Figure 8)**. Other mathematical institutes such as the Mathematical Biology Institute (MBI) at Ohio State University^7^, the Statistical and Applied Mathematical Sciences Institute (SAMSI) in Research Triangle Park^8^, the Institute for Mathematics and Its Applications at University of Minnesota^9^, and the Centre for Plant Integrative Biology at the University of Nottingham^10^ have also hosted workshops for mathematical and quantitative biologists from the undergraduate student to the faculty level.

**Figure 8:**
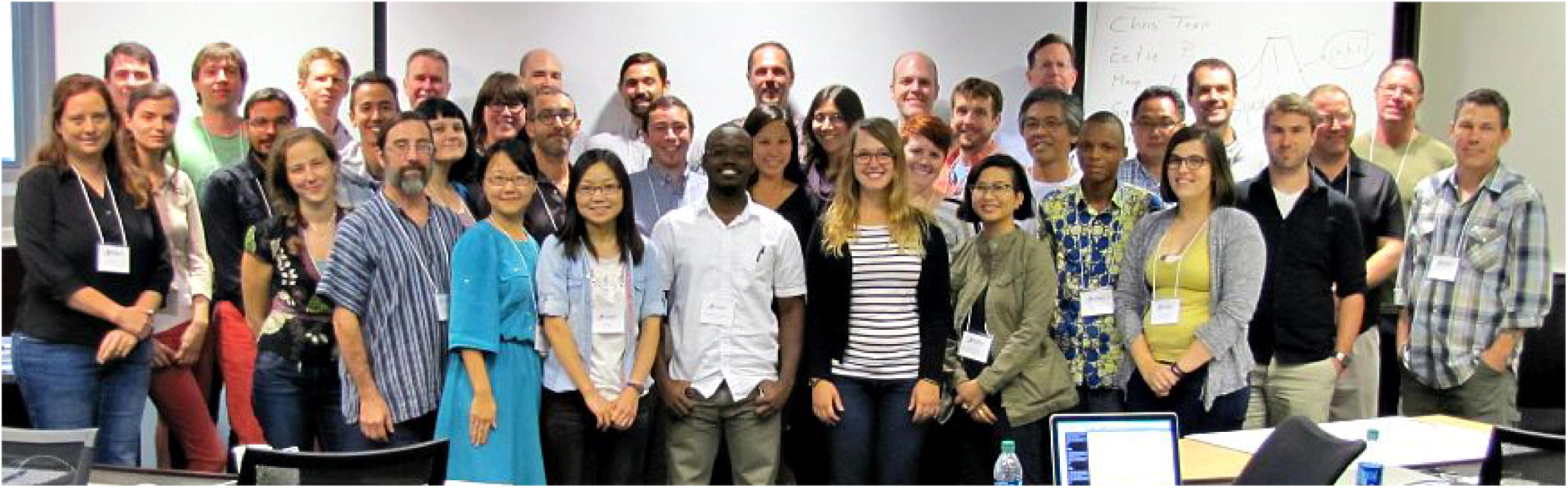
Milestones to accelerate the infusion of math into the plant sciences. Group photo of the authors from the National Institute for Mathematical and Biological Synthesis (NIMBioS) meeting on plant morphological models (University of Tennessee, Knoxville, September 2-4, 2015) that inspired this manuscript. Workshops such as these, bringing mathematicians and plant biologists together, will be necessary to create a new synthesis of plant morphology.

There are efforts to unite biologists and mathematics through initiatives brought forth from The National Science Foundation, including Mathematical Biology Programs^11^ and the Joint DMS/NIGMS Initiative to Support Research at the Interface of the Biological and Mathematical Sciences^12^ (DMS/NIGMS). Outside of the Mathematics and Life Sciences Divisions, the Division of Physics houses a program on the Physics of Living Systems. Societies such as The Society for Mathematical Biology and the Society for Industrial and Applied Mathematics (SIAM) Life Science Activity Group^13^ are focused on the dissemination of research at the intersection of math and biology, creating many opportunities to present research and provide funding. We emphasize the importance that funding opportunities have had and will continue to have in the advancement of plant morphological modeling.

### D. Open Science

Ultimately, mathematicians, computational scientists, and plant biology must unite at the level of jointly collecting data, analyzing it, and doing science together. Open and timely data sharing to benchmark code is a first step to unite these disciplines along with building professional interfaces to bridge between the disciplines (Bucksch et al, 2016).

A number of platforms provide open, public access to datasets, figures, and code that can be shared, including Dryad^14^, Dataverse^15^, and Figshare^16^. Beyond the ability to share data is the question of open data formats and accessibility. For example, in remote sensing research it is unfortunately common that proprietary data formats are used, which prevents their use without specific software. This severely limits the utility and community building aspects of plant morphological research. Beyond datasets, making code openly available, citable, and user-friendly is a means to share methods to analyze data. Places to easily share code include web-based version controlled platforms like Bitbucket^17^ or Github^18^ and software repositories like Sourceforge^19^.

Meta-analysis datasets provide curated resources where numerous published and unpublished datasets related to a specific problem (or many problems) can be accessed by researchers^20^. The crucial element is that data is somehow reflective of universal plant morphological features, bridging the gap between programming languages and biology, as seen in the Root System Markup Language (Lobet et al., 2015) and OpenAlea (Pradal et al., 2008). Bisque is a versatile platform to store, organize, and analyze image data, providing simultaneously open access to data and analyses as well as the requisite computation (Kvilekval et al., 2010). CyVerse^21^ (formerly iPlant) is a similar platform, on which academic users get 100 GB storage for free and can create analysis pipelines that can be shared and reused (Goff et al., 2011). For example, DIRT^22^ is an automatic, high throughput computing platform (Bucksch et al., 2014c; Das et al., 2015) that the public can use hosted on CyVerse using the Texas Advanced Computing Center^23^ (TACC) resources at UT Austin that robustly extracts root traits from digital images. We emphasize the importance of adopting open science policies at the individual investigator and journal level to continue expanding the field of mathematical biology.

## IV. Conclusion: Unleashing geometric and topological potential within the plant sciences

The plant form is inherently morphological, from the shapes of leaves to the hierarchies of branching patterns in shoots and roots. Plant morphology has served as an inspiration for mathematicians to apply new methods to quantify and model the plant form as a result of evolutionary, developmental, and environmental responses **(Figures 1–2)**. Plant morphology is an unresolved mystery to plant biologists, who seek to understand the molecular mechanisms by which such predetermined, yet seemingly endless, variations of organizational patterns emerge.

Never have the resources to study plant morphology been more plentiful. Burgeoning imaging technologies—innovative confocal microscopy, laser ablation tomography, X-ray imaging, MRI, radar, terrestrial laser scanning, among many others—have made detailed 3D models of plants feasible **(Figures 3–4)**. Interest in the hidden half of plant morphology—the root system—has only recently seen a renaissance with technologies capable of penetrating soil and visualizing roots *in situ* **(Figure 5)**.

Integrating observations at different scales is a persistent challenge, such as shoot apical meristem development or the movement of leaves within a tree canopy **(Figures 6–7)**. Modifying plant morphology through molecular biology and breeding is key to develop agricultural outputs and sustainability. Monitoring the morphology of plants in response to a shifting environment is necessary to model global responses to climate change. Cross-disciplinary training of scientists, citizen science, and open science are all necessary components to address these needs (Figure 8). Unleashing the potential of geometric and topological approaches in the plant sciences promises to transform our understanding of both plants and mathematics, and to meet the challenges posed by a future with dwindling and uncertain natural resources.

## Acknowledgements

The authors are grateful to the National Institute for Mathematical and Biological Synthesis (NIMBioS, University of Tennessee, Knoxville) for hosting and funding the workshop “Morphological Plant Modeling: Unleashing geometric and topological potential within the plant sciences” that inspired this manuscript. We thank the reviewers Evelyne Costes and Leo Marcelis for creative and open discussions.

## Footnotes

1 For example, see the White Paper on Citizen Science for Europe, http://www.socientize.eu/sites/default/files/white-paper_0.pdf (retrieved May 29, 2016)

2 List of herbaria, https://en.wikipedia.org/wiki/List_of_herbaria (retrieved May 29, 2016)

3 https://www.usanpn.org/# (retrieved May 29, 2016)

4 http://earthwatch.org/scientific-research/special-initiatives/urban-resiliency (retrieved May 29, 2016)

5 http://www.mytreetracker.org/cwis438/websites/MyTreeTracker/About.php?WebSiteID=23 (retrieved May 29, 2016)

6 http://www.nimbios.org/workshops/WS_plantmorph (retrieved May 29, 2016)

7 https://mbi.osu.edu/ (retrieved May 29, 2016)

8 http://www.samsi.info/ (retrieved May 29, 2016)

9 https://www.ima.umn.edu/ (retrieved May 29, 2016)

10 https://www.cpib.ac.uk/outreach/cpib-summer-school/ (retrieved May 29, 2016)

11 https://www.nsf.gov/funding/pgm_summ.jsp?pims_id=5690 (retrieved May 29, 2016)

12 http://www.nsf.gov/funding/pgm_summ.jsp?pims_id=5300&org=DMS (retrieved May 29, 2016)

13 https://www.siam.org/activity/life-sciences/ (retrieved May 29, 2016)

14 http://datadryad.org/ (retrieved May 29, 2016)

15 http://dataverse.org/ (retrieved May 29, 2016)

16 https://figshare.com/ (retrieved May 29, 2016)

17 https://bitbucket.org/ (retrieved May 29, 2016)

18 https://github.com/ (retrieved May 29, 2016)

19 https://sourceforge.net/ (retrieved May 29, 2016)

20 BAAD: a Biomass And Allometry Database for woody plants, https://github.com/dfalster/baad (retrieved May 29, 2016)

21 http://www.cyverse.org/ (retrieved August 20, 2016)

22 http://dirt.iplantcollaborative.org/ (retrieved August 20, 2016)

23 https://www.tacc.utexas.edu/ (retrieved August 20, 2016)

## References

Aiteanu, F. and Klein, R. (2014). Hybrid tree reconstruction from inhomogeneous point clouds. Visual Comput. 30, 763–771. doi: 10.1007/s00371-014-0977-7.

Bailey, I.W. and Sinnott, E.W. (1915). A botanical index of Cretaceous and Tertiary climates. Science 41, 831–834.

Baldazzi, V., Bertin, N., De Jong, H. and Génard, M. (2012). Towards multiscale plant models: integrating cellular networks. Trends Plant Sci. 17, 728–736. doi: http://dx.doi.org/10.1016/j.tplants.2012.06.012

Band, L.R. and King, J.R. (2012). Multiscale modelling of auxin transport in the plant-root elongation zone. J. Math. Biol. 65, 743–785. doi: 10.1007/s00285-011-0472-y

Band, L.R., Fozard, J.A., Godin, C., Jensen, O.E., Pridmore, T., Bennett, M.J. and King, J.R. (2012). Multiscale systems analysis of root growth and development: modeling beyond the network and cellular scales. Plant Cell 24, 3892–3906. doi:http://dx.doi.org/10.1105/tpc.112.101550

Bao, Y., Aggarwal, P., Robbins, N.E., Sturrock, C.J., Thompson, M.C., Tan, H.Q., Tham, C., Duan, L., Rodriguez, P.L., Vernoux, T. and Mooney, S.J. (2014). Plant roots use a patterning mechanism to position lateral root branches toward available water. Proc. Natl. Acad. Sci. U.S.A. 111, 9319–9324. doi: 10.1073/pnas.1400966111

Barthélémy, D. and Caraglio, Y. (2007). Plant Architecture: A dynamic, multilevel and comprehensive approach to plant form, structure, and ontogeny. Ann. Bot. 99, 375–407. doi: 10.1093/aob/mcl260

Battogtokh D. and Tyson J. J. (2016). A bistable switch mechanism for stem cell domain nucleation in the shoot apical meristem. Front. Plant Sci. 7, 674. doi: 10.3389/fpls.2016.00674

Benfey, P.N. and Mitchell-Olds, T. (2008). From genotype to phenotype: systems biology meets natural variation. Science 320, 495–497. doi: 10.1126/science.1153716

Benz, B.W. and Martin, C.E. (2006). Foliar trichomes, boundary layers, and gas exchange in 12 species of epiphytic Tillandsia (Bromeliaceae). J. Plant Physiol. 163, 648–656. doi: 10.1016/j.jplph.2005.05.008

Bernasconi, G.P. (1994). Reaction-diffusion model for phyllotaxis. Physica D: Nonlinear Phenom. 70, 90–99. doi: 10.1016/0167-2789(94)90058-2

Bookstein, F. L. (1997). Morphometric tools for landmark data: geometry and biology. Cambridge University Press.

Bradshaw, A.D. (1965). Evolutionary significance of phenotypic plasticity in plants. Adv. Genet. 13, 115–155.

Braybrook, S.A. and Jönsson, H. (2016). Shifting foundations: the mechanical cell wall and development. Curr. Opin. Plant Biol. 29, 115–120. doi: http://dx.doi.org/10.1016/j.pbi.2015.12.009

Brooks, T.L.D., Miller, N.D. and Spalding, E.P. (2010). Plasticity of Arabidopsis root gravitropism throughout a multidimensional condition space quantified by automated image analysis. Plant Physiol. 152, 206–216.

Bucksch, A., Lindenbergh, R. and Menenti, M. (2010). SkelTre. Visual Comput. 26, 1283–1300. doi: http://dx.doi.org/10.1007/s00371-010-0520-4

Bucksch, A.K. (2011). Revealing the skeleton from imperfect point clouds. TU Delft, Delft University of Technology.

Bucksch, A. and Fleck, S. (2011). Automated detection of branch dimensions in woody skeletons of fruit tree canopies. Photogramm. Eng. Remote Sens. 77, 229–240. doi: 10.14358/PERS.77.3.229

Bucksch, A. (2014a). A practical introduction to skeletons for the plant sciences. Appl. Plant Sci. 2.apps.1400005. doi:10.3732/apps.1400005

Bucksch, A., Turk, G. and Weitz, J. S. (2014b). The fiber walk: A model of tip-driven growth with lateral expansion. PLoS ONE 9: e85585. doi: 10.1371/journal.pone.0085585

Bucksch, A., Burridge, J., York, L.M., Das, A., Nord, E., Weitz, J.S. and Lynch, J.P. (2014c). Image-based high-throughput field phenotyping of crop roots. Plant Physiol. 166, 470–486. doi: 10.1104/pp.114.243519.

Bucksch, A., Das, A., Schneider, H., Merchant, N., & Weitz, J. S. (2016). Overcoming the Law of the Hidden in Cyberinfrastructures. Trends in Plant Science (online first). doi:http://dx.doi.org/10.1016/j.tplants.2016.11.014

Chickarmane V., Roeder A.H., Tarr P.T., Cunha A., Tobin C., Meyerowitz E. M. (2010). Computational morphodynamics: a modeling framework to understand plant growth. Annu Rev Plant Biol. 2010;61:65–87. doi: 10.1146

Chimungu, J.G., Maliro, M.F., Nalivata, P.C., Kanyama-Phiri, G., Brown, K.M. and Lynch, J.P. (2015). Utility of root cortical aerenchyma under water limited conditions in tropical maize (Zea mays L.). Field Crops Res. 171, 86–98. doi: http://dx.doi.org/10.1016/j.fcr.2014.10.009

Chitwood, D.H., Headland, L.R., Ranjan, A., Martinez, C.C., Braybrook, S.A., Koenig, D.P., Kuhlemeier, C., Smith, R.S. and Sinha, N.R. (2012a). Leaf asymmetry as a developmental constraint imposed by auxin-dependent phyllotactic patterning. Plant Cell 24, 2318–2327. doi: 10.1105/tpc.112.098798

Chitwood, D.H., Naylor, D.T., Thammapichai, P., Weeger, A.C., Headland, L.R. and Sinha, N.R. (2012b). Conflict between intrinsic leaf asymmetry and phyllotaxis in the resupinate leaves of Alstroemeria psittacina. *Front*. Plant Sci. 3:182. doi: 10.3389/fpls.2012.00182

Chitwood, D.H., Kumar, R., Headland, L.R., Ranjan, A., Covington, M.F., Ichihashi, Y., Fulop, D., Jiménez-Gómez, J.M., Peng, J., Maloof, J.N. and Sinha, N.R. (2013). A quantitative genetic basis for leaf morphology in a set of precisely defined tomato introgression lines. Plant Cell 25, 2465–2481. doi: http://dx.doi.org/10.1105/tpc.113.112391

Chitwood, D.H., Ranjan, A., Kumar, R., Ichihashi, Y., Zumstein, K., Headland, L.R., Ostria-Gallardo, E., Aguilar-Martínez, J.A., Bush, S., Carriedo, L. and Fulop, D. (2014a). Resolving distinct genetic regulators of tomato leaf shape within a heteroblastic and ontogenetic context. Plant Cell 26, 3616–3629. doi: http://dx.doi.org/10.1105/tpc.114.130112

Chitwood, D.H., Ranjan, A., Martinez, C.C., Headland, L.R., Thiem, T., Kumar, R., Covington, M.F., Hatcher, T., Naylor, D.T., Zimmerman, S. and Downs, N. (2014b). A modern ampelography: a genetic basis for leaf shape and venation patterning in grape. Plant Physiol. 164, 259–272. doi: http://dx.doi.org/10.1104/pp.113.229708

Chitwood, D.H. and Topp, C.N. (2015). Revealing plant cryptotypes: defining meaningful phenotypes among infinite traits. Curr. Opin. Plant Biol. 24, 54–60. doi: http://dx.doi.org/10.1016/j.pbi.2015.01.009

Chitwood, D.H., Rundell, S.M., Li, D.Y., Woodford, Q.L., Tommy, T.Y., Lopez, J.R., Greenblatt, D., Kang, J. and Londo, J.P. (2016). Climate and developmental plasticity: interannual variability in grapevine leaf morphology. Plant Physiol. 170, 1480–91. doi: http://dx.doi.org/10.1104/pp.15.01825

Clark, R.M., Wagler, T.N., Quijada, P. and Doebley, J. (2006). A distant upstream enhancer at the maize domestication gene tb1 has pleiotropic effects on plant and inflorescent architecture. Nat. Genet. 38, 594–597. doi: 10.1038/ng1784

Clark, R.T., MacCurdy, R.B., Jung, J.K., Shaff, J.E., McCouch, S.R., Aneshansley, D.J. and Kochian, L.V. (2011). Three-dimensional root phenotyping with a novel imaging and software platform. Plant Physiol. 156, 455–465. doi: 10.1104/pp.110.169102

Clausen, J., Keck, D.D. and Hiesey, W.M. (1941). Regional differentiation in plant species. Am. Nat. 75, 231–250.

Cohen, J.E. (2004). Mathematics is biology’s next microscope, only better; biology is mathematics’ next physics, only better. PLoS Biol. 2: e439. doi:10.1371/journal.pbio.0020439

Cordell, S., Goldstein, G., Mueller-Dombois, D., Webb, D. and Vitousek, P.M. (1998). Physiological and morphological variation in Metrosideros polymorpha, a dominant Hawaiian tree species, along an altitudinal gradient: the role of phenotypic plasticity. Oecologia 113, 188–196. doi:10.1007/s004420050367

Corney, D., Clark, J.Y., Tang, H.L. and Wilkin, P. (2012). Automatic extraction of leaf characters from herbarium specimens. Taxon 61, 231–244.

Danjon, F., Bert, D., Godin, C., & Trichet, P. (1999). Structural root architecture of 5-year-old Pinus pinaster measured by 3D digitising and analysed with AMAPmod. Plant and Soil, 217(1–2), 49–63.

Das, A., Schneider, H., Burridge, J., Ascanio, A.K.M., Wojciechowski, T., Topp, C.N., Lynch, J.P., Weitz, J.S. and Bucksch, A. (2015). Digital imaging of root traits (DIRT): a high-throughput computing and collaboration platform for field-based root phenomics. Plant Methods 11:51. doi: 10.1186/s13007-015-0093-3

DeWitt T.J. (2016). Expanding the phenotypic plasticity paradigm to broader views of trait space and ecological function. Curr. Zool. 62, 463–473. doi:http://dx.doi.org/10.1093/cz/zow085

DeWitt T.J. Scheiner S.M. eds. (2004). Phenotypic plasticity: functional and conceptual approaches (No. 576.53 P44). Oxford: Oxford University Press.

Díaz, S., Kattge, J., Cornelissen, J.H., Wright, I.J., Lavorel, S., Dray, S., Reu, B., Kleyer, M., Wirth, C., Prentice, I.C., Garnier, E. et al. (2016). The global spectrum of plant form and function. Nature 529, 167–171. doi:10.1038/nature16489

Doebley, J. (2004). The genetics of maize evolution. Annu. Rev. Genet. 38, 37–59. doi:10.1146/annurev.genet.38.072902.092425

Douady, S. and Coudert, Y. (1996). Phyllotaxis as a dynamical self organizing process part I: the spiral modes resulting from time-periodic iterations. J. Theoret. Biol. 178, 255–273. doi:10.1006/jtbi.1996.0024

Drew, M.C. (1975). Comparison of the effects of a localised supply of phosphate, nitrate, ammonium and potassium on the growth of the seminal root system, and the shoot, in barley. New Phytol. 75, 479–490. doi: 10.1111/j.1469-8137.1975.tb01409.x

Dunbabin, V. M., Postma, J. A., Schnepf, A., Pagès, L., Javaux, M., Wu, L., … and Diggle, A. J. (2013). Modelling root–soil interactions using three–dimensional models of root growth, architecture and function. Plant Soil 372, 93–124. doi:10.1007/s11104-013-1769-y

Edelsbrunner, H. and Harer, J. (2010). Computational topology: an introduction. Rhode Island: American Mathematical Society.

Ennos, A.R. (1997). Wind as an ecological factor. Trends Ecol. Evol. 12, 108–111. doi:10.1016/S0169-5347(96)10066-5

Esau, K. (1960). Anatomy of Seed Plants. New York: John Wiley & Sons Inc.

Etnier, S.A. and Vogel, S. (2000). Reorientation of daffodil (Narcissus: Amaryllidaceae) flowers inwind: drag reduction andtorsional flexibility. Am. J. Bot. 87, 29–32.

Feng, Z., Chen, Y., Hakala, T., Hyyppä, J. (2016). Range Calibration of Airborne Profiling Radar Used in Forest Inventory. IEEE Geoscience and Remote Sensing Society

Fiorani, F., Rascher, U., Jahnke, S. and Schurr, U. (2012). Imaging plants dynamics in heterogenic environments. Curr. Opin. Biotechnol. 23, 227–235. doi:/10.1016/j.copbio.2011.12.010

Fisher, R.A. (1925). Statistical methods for research workers. Guilford: Genesis Publishing Pvt Ltd.

Fisher, R.A. (1936). The use of multiple measurements in taxonomic problems. Ann. Eugenics 7, 179–188.

Fitter, A. H. (1987). An architectural approach to the comparative ecology of plant root systems. New phytologist, 106(s1), 61–77.

Frary, A., Nesbitt, T.C., Frary, A., Grandillo, S., Van Der Knaap, E., Cong, B., Liu, J., Meller, J., Elber, R., Alpert, K.B. and Tanksley, S.D. (2000). fw2. 2: a quantitative trait locus key to the evolution of tomato fruit size. Science 289, 85–88. doi: 10.1126/science.289.5476.85

Friedman, W.E. and Diggle, P.K. (2011). Charles Darwin and the origins of plant evolutionary developmental biology. Plant Cell 23, 1194–1207. doi:http://dx.doi.org/10.1105/tpc.111.084244

Fujita, H., Toyokura, K., Okada, K. and Kawaguchi, M. (2011). Reaction-diffusion pattern in shoot apical meristem of plants. PLoS ONE 6: p.e18243. doi:http://dx.doi.org/10.1371/journal.pone.0018243

Galkovskyi, T., Mileyko, Y., Bucksch, A., Moore, B., Symonova, O., Price, C. A., … & Harer, J. (2012). GiA Roots: software for the high throughput analysis of plant root system architecture. BMC Plant Biol. 12: 116. doi: 10.1186/1471-2229-12-116

Godin, C., Costes, E., & Sinoquet, H. (1999). A method for describing plant architecture which integrates topology and geometry. Annals of botany, 84(3), 343–357.

Godin, C. and Ferraro, P. (2010). Quantifying the degree of self-nestedness of trees: Application to the structural analysis of plants. IEEE/ACM Trans. Comput. Biol. Bioinform. 7, 688–703. doi:10.1109/TCBB.2009.29

Goethe, J.W. (1790). Versuch die Metamorphose der Pflanzen zu erklaren. Gotha: Carl Wilhelm Ettinger.

Goff, S.A., Vaughn, M., McKay, S., Lyons, E., Stapleton, A.E., Gessler, D., Matasci, N., Wang, L., Hanlon, M., Lenards, A. and Muir, A. (2011). The iPlant collaborative: cyberinfrastructure for plant biology. *Front*. Plant Sci. 2:34. doi: 10.3389/fpls.2011.00034

Gordon, S. P., Chickarmane V.S., Ohno C., Meyerowitz E.M. (2009). Multiple feedback loops through cytokinin signaling control stem cell number within the Arabidopsis shoot meristem. PNAS U S A. 2009 Sep 22;106(38):16529–34.

Grant, R.H. (1983). The scaling of flow in vegetative structures. Boundary-Layer Meteorol. 27, 171–184. doi: 10.1007/BF00239613

Green, P.B. (1999). Expression of pattern in plants: combining molecular and calculus-based biophysical paradigms. Am. J. Bot. 86, 1059–1076.

Gupta, M.D. and Nath, U. (2015). Divergence in patterns of leaf growth polarity is associated with the expression divergence of miR396. Plant Cell 27, 2785–2799. doi:http://dx.doi.org/10.1105/tpc.15.00196

Gutenkunst, R.N., Waterfall, J.J., Casey, F.P., Brown, K.S., Myers, C.R. and Sethna, J.P. (2007). Universally sloppy parameter sensitivities in systems biology models. PLoS Comput. Biol. 3:e189. doi: 10.1371/journal.pcbi.0030189

Hallé, F. (1971). Architecture and growth of tropical trees exemplified by the Euphorbiaceae. Biotropica 3, 56–62.

Hallé, F. (1986). Modular Growth in Seed Plants. Philosophical Transactions of the Royal Society B 313, 77–87

Hamant, O., Heisler, M.G., Jönsson, H., Krupinski, P., Uyttewaal, M., Bokov, P., Corson, F., Sahlin, P., Boudaoud, A., Meyerowitz, E.M. and Couder, Y. (2008). Developmental patterning by mechanical signals in Arabidopsis. Science 322, 1650–1655. doi:10.1126/science.1165594

Hofhuis, H., Moulton, D., Lessinnes, T., Routier-Kierzkowska, A.L., Bomphrey, R.J., Mosca, G., Reinhardt, H., Sarchet, P., Gan, X., Tsiantis, M., Ventikos, Y., Walker, S., Goriely, A., Smith, R., Hay, A. (2016). Morphomechanical innovation drives explosive seed dispersal. Cell 166, 222–33. doi: 10.1016/j.cell.2016.05.002

Hohm, T., Zitzler, E. and Simon, R. (2010). A dynamic model for stem cell homeostasis and patterning in Arabidopsis meristems. PLoS ONE 5:e9189. doi:http://dx.doi.org/10.1371/journal.pone.0009189

Horn, H.S. (1971). The adaptive geometry of trees (Vol. 3). Princeton University Press.

Jensen, O.E. and Fozard, J.A. (2015). Multiscale models in the biomechanics of plant growth. Physiology 30, 159–166. doi: 10.1152/physiol.00030.2014

Kaasalainen, S., Holopainen, M., Karjalainen, M., Vastaranta, M., Kankare, V., Karila, K. and Osmanoglu, B. (2015). Combining lidar and synthetic aperture radar data to estimate forest biomass: status and prospects. Forests 6, 252–270. doi:10.3390/f6010252

Kaplan, D.R. (2001). The science of plant morphology: definition, history, and role in modern biology. Am. J. Bot. 88, 1711–1741.

Kendall, D.G. (1984). Shape manifolds, procrustean metrics, and complex projective spaces. Bull. Lond. Math. Soc. 16, 81–121. doi: 10.1112/blms/16.2.81

Kenis, K., and Keulemans, J. (2007). Study of tree architecture of apple (Malus× domestica Borkh.) by QTL analysis of growth traits. Molecular Breeding, 19(3), 193–208.

Kimura, S., Koenig, D., Kang, J., Yoong, F.Y. and Sinha, N. (2008). Natural variation in leaf morphology results from mutation of a novel KNOX gene. Curr. Biol. 18, 672–677. doi: 10.1016/j.cub.2008.04.008

Kitazawa, M.S. and Fujimoto, K. (2015). A Dynamical Phyllotaxis Model to Determine Floral Organ Number. PLoS Comput. Biol. 11: p.e1004145. doi:10.1371/journal.pcbi.1004145

Ku, L.X., Zhao, W.M., Zhang, J., Wu, L.C., Wang, C.L., Wang, P.A., Zhang, W.Q. and Chen, Y.H. (2010). Quantitative trait loci mapping of leaf angle and leaf orientation value in maize (*Zea mays* L.). Theoret. Appl. Genet. 121, 951–959. doi:10.1007/s00122-010-1364-z

Kuhl, F.P. and Giardina, C.R. (1982). Elliptic Fourier features of a closed contour. Comput. Graph. Image Process. 18, 236–258. doi:10.1016/0146-664X(82)90034-X

Kumi, F., Hanping, M., Jianping, H. and Ullah, I. (2015). Review of applying X-ray computed tomography for imaging soil-root physical and biological processes. Int. J. Agric. Biol. Eng. 8, 1–14.

Kurth, W. (1994). Growth grammar interpreter grogra 2.4-a software tool for the 3-dimensional interpretation of stochastic, sensitive growth grammars in the context of plant modelling. Göttingen : Forschungszentrum Waldökosysteme der Universität Göttingen.

Kvilekval, K., Fedorov, D., Obara, B., Singh, A. and Manjunath, B.S. (2010). Bisque: a platform for bioimage analysis and management. Bioinformatics 26, 544–552. doi: 10.1093/bioinformatics/btp699

Laga, H., Kurtek, S., Srivastava, A. and Miklavcic, S. J. (2014). Landmark-free statistical analysis of the shape of plant leaves. *J. Theoret*. Biol. 363, 41–52. doi: 10.1016/j.jtbi.2014.07.036

Langlade, N.B., Feng, X., Dransfield, T., Copsey, L., Hanna, A.I., Thébaud, C., Bangham, A., Hudson, A. and Coen, E. (2005). Evolution through genetically controlled allometry space. Proc. Natl. Acad. Sci. U.S.A. 102, 10221–10226. doi: 10.1073/pnas.0504210102

Leiboff, S., Li, X., Hu, H.C., Todt, N., Yang, J., Li, X., Yu, X., Muehlbauer, G.J., Timmermans, M.C., Yu, J. and Schnable, P.S. (2015). Genetic control of morphometric diversity in the maize shoot apical meristem. Nat. Commun. 6:8974. doi: 10.1038/ncomms9974

Lobet, G., Pagès, L., and Draye, X. (2011). A novel image-analysis toolbox enabling quantitative analysis of root system architecture. Plant physiol., 157, 29–39.doi:http://dx.doi.org/10.1104/pp.111.179895

Lobet, G., Pound, M.P., Diener, J., Pradal, C., Draye, X., Godin, C., Javaux, M., Leitner, D., Meunier, F., Nacry, P. and Pridmore, T.P. (2015). Root system markup language: toward a unified root architecture description language. Plant Physiol. 167, 617–627. doi: 10.1104/pp.114.253625

Lynch, J. P., & Brown, K. M. (2012). New roots for agriculture: exploiting the root phenome. Philos. Trans. R. Soc. B 367, 1598–1604. doi:10.1098/rstb.2011.0243

Lynch, J.P. (2013). Steep, cheap and deep: an ideotype to optimize water and N acquisition by maize root systems. Ann. Bot. 112, 347–357. doi:10.1093/aob/mcs293

Lynch, J.P. (2015). Root phenes that reduce the metabolic costs of soil exploration: opportunities for 21st century agriculture. Plant Cell Environ. 38, 1775–1784. doi: 10.1111/pce.12451

MacPherson, R. and Schweinhart, B. (2012). Measuring shape with topology. J. Math. Phys. 53: 073516. doi: 10.1063/1.4737391

Martinez, C.C., Chitwood, D.H., Smith, R.S. and Sinha, N.R. (2016). Left-right leaf asymmetry in decussate and distichous phyllotactic systems. bioRxiv, p.043869. doi.http://dx.doi.org/10.1101/043869

Mayr, E. (1981). Biological classification: toward a synthesis of opposing methodologies. Science 214, 510–516. doi:10.1126/science.214.4520.510

Meijón, M., Satbhai, S.B., Tsuchimatsu, T. and Busch, W. (2014). Genome-wide association study using cellular traits identifies a new regulator of root development in Arabidopsis. Nat. Genet. 46, 77–81. doi: 10.1038/ng.2824

Meinhardt, H. and Gierer, A. (1974). Applications of a theory of biological pattern formation based on lateral inhibition. J. Cell Sci. 15, 321–346.

Meinhardt, H. (1976). Morphogenesis of lines and nets. Differentiation 6, 117–123. doi: 10.1111/j.1432-0436.1976.tb01478.x

Meinhardt, H. (2004). Out-of-phase oscillations and traveling waves with unusual properties: the use of three-component systems in biology. Physica D: Nonlinear Phenomena 199, 264–277. doi: http://dx.doi.org/10.1016/j.physd.2004.08.018

Miller, N.D., Parks, B.M. and Spalding, E.P. (2007). Computer-vision analysis of seedling responses to light and gravity. Plant J. 52, 374–381. doi: 10.1111/j.1365- 313X.2007.03237.x

Milnor, J.W. (1963). Morse theory. Princeton: Princeton University Press.

Moulia, B., & Fournier, M. (2009). The power and control of gravitropic movements in plants: a biomechanical and systems biology view. Journal of experimental botany, 60(2), 461–486.

Monforte, A.J., Diaz, A.I., Caño-Delgado, A. and van der Knaap, E. (2014). The genetic basis of fruit morphology in horticultural crops: lessons from tomato and melon. J. Exp. Bot. 65, 4625–4637. doi: 10.1093/jxb/eru017.

Nicotra, A.B., Atkin, O.K., Bonser, S.P., Davidson, A.M., Finnegan, E.J., Mathesius, U., Poot, P., Purugganan, M.D., Richards, C.L., Valladares, F. and van Kleunen, M. (2010). Plant phenotypic plasticity in a changing climate. Trends Plant Sci. 15, 684–692. doi: http://dx.doi.org/10.1016/j.tplants.2010.09.008.

Nielsen, K. L., Lynch, J. P., Jablokow, A. G. and Curtis, P. S. (1994). Carbon cost of root systems: an architectural approach. In Belowground Responses to Rising Atmospheric CO2: Implications for Plants, Soil Biota, and Ecosystem Processes (pp. 161–169). Springer Netherlands.

Niklas, K.J. (1992). Plant biomechanics: an engineering approach to plant form and function. Chicago: University of Chicago Press.

Niklas, K.J. (1994). Plant allometry: the scaling of form and process. Chicago: University of Chicago Press.

Niklas, K.J. (1997). The evolutionary biology of plants. Chicago: University of Chicago Press.

Palubicki, W., Horel, K., Longay, S., Runions, A., Lane, B., Měch, R., & Prusinkiewicz, P. (2009). Self-organizing tree models for image synthesis. ACM Transactions on Graphics (TOG), 28(3), 58.

Palubicki, W. (2013). A Computational Study of Tree Architecture (Doctoral dissertation, University of Calgary).

Paran, I. and van der Knaap, E. (2007). Genetic and molecular regulation of fruit and plant domestication traits in tomato and pepper. J. Exp. Bot. 58, 3841–3852. doi: 10.1093/jxb/erm257

Peaucelle, A., Braybrook, S.A., Le Guillou, L., Bron, E., Kuhlemeier, C. and Höfte, H. (2011). Pectin-induced changes in cell wall mechanics underlie organ initiation in Arabidopsis. Curr. Biol. 21, 1720–1726. doi: 10.1016/j.cub.2011.08.057

Postma, J.A. and Lynch, J.P. (2011). Root cortical aerenchyma enhances the growth of maize on soils with suboptimal availability of nitrogen, phosphorus, and potassium. Plant Physiol. 156, 1190–1201. doi: 10.1104/pp.111.175489

Pradal, C., Dufour-Kowalski, S., Boudon, F., Fournier, C. and Godin, C. (2008). OpenAlea: a visual programming and component-based software platform for plant modelling. Funct. Plant Biol. 35, 751–760. doi: 10.1071/FP08084

Prusinkiewicz, P., Mündermann, L., Karwowski, R. and Lane, B. (2001). The use of positional information in the modeling of plants. In Proceedings of the 28th annual conference on Computer graphics and interactive techniques (289-300). ACM.

Prusinkiewicz, P. (2004). “Self-similarity in plants: Integrating mathematical and biological perspectives”, in Thinking in Patterns: Fractals and Related Phenomena in Nature, ed. M. Novak (Singapore: World Scientific), 103–118.

Prusinkiewicz, P. and de Reuille, P.B. (2010). Constraints of space in plant development. J. Exp. Bot. 61, 2117–2129. doi:10.1093/jxb/erq081

Prusinkiewicz, P. and Lindenmayer, A. (2012). The algorithmic beauty of plants. New York: Springer Science & Business Media.

Raumonen, P., Kaasalainen, M., Åkerblom, M., Kaasalainen, S., Kaartinen, H., Vastaranta, M., Holopainen, M., Disney, M. and Lewis, P. (2013). Fast automatic precision tree models from terrestrial laser scanner data. Remote Sens. 5, 491–520. doi:10.3390/rs5020491

Razak, K. A., Bucksch, A., Damen, M., van Westen, C., Straatsma, M., & de Jong, S. (2013). Characterizing tree growth anomaly induced by landslides using LiDAR. In Landslide Science and Practice (pp. 235–241). Springer Berlin Heidelberg.

Rellán-Álvarez, R., Lobet, G., Lindner, H., Pradier, P.L., Sebastian, J., Yee, M.C., Geng, Y., Trontin, C., LaRue, T., Schrager-Lavelle, A. and Haney, C.H. (2015). GLO-Roots: an imaging platform enabling multidimensional characterization of soil-grown root systems. Elife 4: e07597. doi: 10.7554/eLife.07597

Rellán-Álvarez, R., Lobet, G. and Dinneny, J.R. (2016). Environmental Control of Root System Biology. Annu. Rev. Plant Biol. 67, 619–642. doi: 10.1146/annurev-arplant-043015-111848

Remmler, L. and Rolland-Lagan, A.G. (2012). Computational method for quantifying growth patterns at the adaxial leaf surface in three dimensions. Plant Physiol. 159, 27–39. doi:10.1104/pp.112.194662

Reuter, M., Biasotti, S., Giorgi, D., Patanè, G. and Spagnuolo, M. (2009). Discrete Laplace– Beltrami operators for shape analysis and segmentation. Comput. Graph. 33, 381–390. doi:http://dx.doi.org/10.1016/j.cag.2009.03.005

Robbins, N.E. and Dinneny, J.R. (2015). The divining root: moisture-driven responses of roots at the micro-and macro-scale. J. Exp. Bot. 66, 2145–2154.doi:10.1093/jxb/eru496

Roden, J.S. and Pearcy, R.W. (1993). Effect of leaf flutter on the light environment of poplars. Oecologia 93, 201–207. doi: 10.1007/BF00317672

Rolland-Lagan, A.G., Bangham, J.A. and Coen, E. (2003). Growth dynamics underlying petal shape and asymmetry. Nature 422, 161–163. doi:10.1038/nature01443

Ron, M., Dorrity, M.W., de Lucas, M., Toal, T., Hernandez, R.I., Little, S.A., Maloof, J.N., Kliebenstein, D.J. and Brady, S.M. (2013). Identification of novel loci regulating interspecific variation in root morphology and cellular development in tomato. Plant Physiol. 162, 755–768. doi: 10.1104/pp.113.217802

Royer, D.L., Meyerson, L.A., Robertson, K.M. and Adams, J.M. (2009). Phenotypic plasticity of leaf shape along a temperature gradient in *Acer rubrum*. PLoS ONE 4: e7653.doi:10.1371/journal.pone.0007653

Runions, A., Lane, B., and Prusinkiewicz, P. (2007). “Modeling Trees with a Space Colonization Algorithm”. NPH, 63–70.

Ryan, D. (2013). The Global Plants Initiative celebrates its achievements and plans for the future. Taxon 62, 417–418. doi: http://dx.doi.org/10.12705/622.26

Sahlin P., Melke P., Jönsson H. (2011). Models of sequestration and receptor cross-talk for explaining multiple mutants in plant stem cell regulation. BMC Syst Biol. 2011 Jan 5;5:2.doi: 10.1186/1752-0509-5-2.

Schreuder, M.D., Brewer, C.A. and Heine, C. (2001). Modelled influences of non-exchanging trichomes on leaf boundary layers and gas exchange. J. Theoret. Biol. 210, 23–32. doi:10.1006/jtbi.2001.2285

Segura, V., Durel, C. E., and Costes, E. (2009). Dissecting apple tree architecture into genetic, ontogenetic and environmental effects: QTL mapping. Tree genetics & genomes, 5(1), 165–179.

Seidel, D., Schall, P., Gille, M. and Ammer, C. (2015). Relationship between tree growth and physical dimensions of Fagus sylvatica crowns assessed from terrestrial laser scanning. iForest. 8, 735–742. doi: 10.3832/ifor1566-008

Shapiro B.E., Meyerowitz E.M., Mjolsness E. (2013). Using cellzilla for plant growth simulations at the cellular level. Front Plant Sci. 2013 Oct 16;4:408.

Silk, W.K. and Erickson, R.O. (1979). Kinematics of plant growth. J. Theoret. Biol. 76, 481–501. doi:10.1016/0022-5193(79)90014-6

Slovak, R., Göschl, C., Su, X., Shimotani, K., Shiina, T. and Busch, W. (2014). A scalable open-source pipeline for large-scale root phenotyping of *Arabidopsis*. Plant Cell 26, 2390–2403. doi: 10.1105/tpc.114.124032

Smith, G.F., Steenkamp, Y., Klopper, R.R., Siebert, S.J. and Arnold, T.H. (2003). The price of collecting life. Nature 422, 375–376. doi:10.1038/422375a

Spalding, E.P. and Miller, N.D. (2013). Image analysis is driving a renaissance in growth measurement. Curr. Opin. Plant Biol. 16, 100–104. doi: http://dx.doi.org/10.1016/j.pbi.2013.01.001

Steeves, T.A. and Sussex, I.M. (1989). Patterns in plant development. New York: Cambridge University Press.

Sudret, B. (2008). Global sensitivity analysis using polynomial chaos expansions. Reliab. Eng. Syst. Saf. 93, 964–979. doi: http://dx.doi.org/10.1016/j.ress.2007.04.002.

Symonova, O., Topp, C. N., & Edelsbrunner, H. (2015). DynamicRoots: a software platform for the reconstruction and analysis of growing plant roots. PLoS ONE, 10: e0127657. doi:10.1371/journal.pone.0127657.

Thom, A.S. (1968). The exchange of momentum, mass, and heat between an artificial leaf and the airflow in a wind-tunnel. Q. J. R. Meteorol. Soc. 94, 44–55. doi: 10.1002/qj.49709439906.

Thompson, A.M., Yu, J., Timmermans, M.C., Schnable, P., Crants, J.C., Scanlon, M.J. and Muehlbauer, G.J. (2015). Diversity of maize shoot apical meristem architecture and its relationship to plant morphology. G3. Genes Genomes Genet. 5, 819–827. doi: 10.1534/g3.115.017541.

Tian, F., Bradbury, P.J., Brown, P.J., Hung, H., Sun, Q., Flint-Garcia, S., Rocheford, T.R., McMullen, M.D., Holland, J.B. and Buckler, E.S. (2011). Genome-wide association study of leaf architecture in the maize nested association mapping population. Nat. Genet. 43, 159–162. doi:10.1038/ng.746.

Tisné, S., Reymond, M., Vile, D., Fabre, J., Dauzat, M., Koornneef, M. and Granier, C. (2008). Combined genetic and modeling approaches reveal that epidermal cell area and number in leaves are controlled by leaf and plant developmental processes in Arabidopsis. Plant Physiol. 148, 1117–1127. doi: http://dx.doi.org/10.1104/pp.108.124271.

Topp, C.N., Iyer-Pascuzzi, A.S., Anderson, J.T., Lee, C.R., Zurek, P.R., Symonova, O., Zheng, Y., Bucksch, A., Mileyko, Y., Galkovskyi, T. and Moore, B.T. (2013). 3D phenotyping and quantitative trait locus mapping identify core regions of the rice genome controlling root architecture. Proc. Natl. Acad. Sci. U.S.A. 110, E1695–E1704. doi: 10.1073/pnas.1304354110.

Truong, S.K., McCormick, R.F., Rooney, W.L. and Mullet, J.E. (2015). Harnessing genetic variation in leaf angle to increase productivity of *Sorghum bicolor*. Genetics 201, 1229–1238. doi: 10.1534/genetics.115.178608.

Turing, A.M. (1952). The chemical basis of morphogenesis. Philos. Trans. R. Soc. B 237, 37–72.

Turing, A. M. (1992). In *[Collected works]; Collected works of AM Turing.[3]*. Morphogenesis. Saunders, P.T. ed., North-Holland.

Uga, Y., Sugimoto, K., Ogawa, S., Rane, J., Ishitani, M., Hara, N., Kitomi, Y., Inukai, Y., Ono, K., Kanno, N. and Inoue, H. (2013). Control of root system architecture by DEEPER ROOTING 1 increases rice yield under drought conditions. Nat. Genet. 45, 1097–1102. doi:10.1038/ng.2725

van Dusschoten, D., Metzner, R., Kochs, J., Postma, J.A., Pflugfelder, D., Buehler, J., Schurr, U. and Jahnke, S. (2016). Quantitative 3D analysis of plant roots growing in soil using Magnetic Resonance Imaging. Plant Physiol. pp.01388.2015. doi:10.1104/pp.15.01388

Via, S. and Lande, R. (1985). Genotype-environment interaction and the evolution of phenotypic plasticity. Evolution 39, 522505–522522.

Vogel, S. (1984). Drag and flexibility in sessile organisms. Am. Zool. 24, 37–44. doi: http://dx.doi.org/10.1093/icb/24.1.37

Vogel, S. (1989). Drag and reconfiguration of broad leaves in high winds. J. Exp. Bot. 40, 941–948. doi: 10.1093/jxb/40.8.941

Vogel, S. (2006). Drag reduction by leaf aquaplaning in Hexastylis (Aristolochiaceae) and other plant species in floods. J. North Am. Benthological Soc. 25, 2–8.

Vogel, S. (2012). The life of a leaf. Chicago: The University of Chicago Press.

Vosselman, G. and Maas, H.G. eds. (2010). Airborne and terrestrial laser scanning. Caithness: Whittles Publishing.

Watanabe, T., Hanan, J. S., Room, P. M., Hasegawa, T., Nakagawa, H., & Takahashi, W. (2005). Rice morphogenesis and plant architecture: measurement, specification and the reconstruction of structural development by 3D architectural modelling. Annals of botany, 95(7), 1131–1143.

Wiens, J.J. (2000). “Coding morphological variation within species and higher taxa for phylogenetic analysis” in Phylogenetic Analysis of Morphological Data, ed J.J. Wiens (Washington, D.C.: Smithsonian Institution Press), 115–145.

Wilf, P., Zhang, S., Chikkerur, S., Little, S.A., Wing, S.L., Serre, T. (2016). Computer vision cracks the leaf code. Proc. Natl. Acad. Sci. U.S.A. 113, 3305–3310. doi: 10.1073/pnas.1524473113

Willis L, Refahi Y., Wightman R., Landrein B., Teles J., Huang K.C., Meyerowitz E.M., Jönsson H. (2016). Cell size and growth regulation in the Arabidopsis thaliana apical stem cell niche. PNAS U S A. 2016 Dec 20;113(51):E8238–E8246. doi: 10.1073/pnas.1616768113.

Woltereck, R. (1909). “Weitere experimentelle Untersuchungen über Artveränderung, speziel über das Wesen quantitativer Artunterschiede bei Daphniden.” Gesellsc("Further investigations of type variation, specifically concerning the nature of quantitative differences between varieties of Daphnia." Verhandlungen der Deutschen zoologischen Gesellschaft 19(1909): 110–73.

Zhang, J., Ku, L.X., Han, Z.P., Guo, S.L., Liu, H.J., Zhang, Z.Z., Cao, L.R., Cui, X.J. and Chen, Y.H. (2014). The ZmCLA4 gene in the qLA4-1 QTL controls leaf angle in maize (Zea mays L.). J. Exp. Bot. 65, 5063–5076. doi: 10.1093/jxb/eru271

Zhu, J., Kaeppler, S. M. and Lynch, J. P. (2005). Mapping of QTLs for lateral root branching and length in maize (Zea mays L.) under differential phosphorus supply. Theor. Appl. Genet. 111, 688–695.

Zhu, J., Brown, K.M. and Lynch, J.P. (2010). Root cortical aerenchyma improves the drought tolerance of maize (Zea mays L.). Plant Cell Environ. 33, 740–749. doi: 10.1111/j.1365-3040.2009.02099.x

Zurek, P.R., Topp, C.N. and Benfey, P.N. (2015). Quantitative trait locus mapping reveals regions of the maize genome controlling root system architecture. Plant Physiol. 167, 1487–1496. doi: http://dx.doi.org/10.1104/pp.114.251751

